# Mindin differentially regulates fibroblast subpopulations via distinct members of the Src family kinases during fibrogenesis

**DOI:** 10.1101/2022.05.17.492347

**Authors:** Sunny Kataria, Isha Rana, Krithika Badarinath, Rania F Zaarour, Gaurav Kansagara, Ravindra K Zirmire, Binita Dam, Pankaj Kumar, Akash Gulyani, Colin Jamora

## Abstract

Fibrosis is the result of excessive deposition of extracellular matrix (ECM) proteins leading to tissue hardening and loss of organ function. A central player driving fibrosis is the activated fibroblast, which exhibits enhanced migration, proliferation, contraction, and ECM production. However, this raises an interesting puzzle of whether the same fibroblast performs all of the processes that fall under the umbrella term of “activation”. Given the heterogeneity of fibroblasts in connective tissues, there are subpopulations of fibroblasts that perform specific functions that are under different regulatory controls. Using a transgenic mouse model of skin fibrosis, we find that the secretion of Mindin from Snail transgenic keratinocytes differentially alters the characteristic of distinct fibroblast subpopulations. Mindin induces migration and inflammatory gene expression of the Sca1+ subpopulation of dermal fibroblasts in a Fyn kinase-dependent manner. On the other hand, Mindin increases the contractile behaviour and collagen production in the papillary CD26+ dermal fibroblasts via c-Src. Moreover, in the context of the fibrotic microenvironment of the tumour stroma, we found that differential responses of resident fibroblasts subpopulations to Mindin extend to the generation of functionally heterogeneous cancer-associated fibroblasts (CAFs). Overall, this work highlights the importance of Mindin in mediating the cellular and signalling heterogeneity of dermal fibroblasts in skin fibrosis and cancer.

## INTRODUCTION

Fibrosis is the leading cause of death in many chronic diseases and the central players in this pathophysiology are fibroblasts. The key events in the pathogenesis can be conceptualised as a chronic exaggeration of wound healing processes that leads to scarring ^1–3^. The processes involved in wound healing can be broadly divided into four overlapping phases – hemostasis, inflammatory phase, proliferative phase, and the remodelling phase ^4^. During the inflammatory phase, the fibroblasts are activated and migrate to the site of injury. The fibroblasts at the wound site proliferate and differentiate into myofibroblasts where they perform a number of functions and coordinate multiple aspects of the wound healing program ^5^. While the number of activated fibroblasts dissipates following normal wound healing, this is not the case with fibrotic diseases which are associated with the persistent presence of activated fibroblasts ^5,6^. However, to understand what causes the persistence of activated fibroblasts in pathological scenarios such as fibrosis, it is important to understand what activates them and how they remain in a chronically active state.

A major complication in understanding fibroblast regulation and function is that fibroblasts in the dermal compartment of the skin are spatially and functionally heterogeneous ^7–12^. Fibroblasts in the neonatal skin arise from two distinct lineages. The upper lineage, marked by CD26+, forms the papillary dermis and contributes to hair follicle generation, arrector pili muscle, and epidermal homeostasis ^8,11^. The lower dermal lineage, marked by Sca1+ contributes to deposition of the majority of fibrous collagen and can differentiate into adipocytes to maintain the skin dermal white adipose layer ^8,11,13^. In wound healing, the Sca1+ fibroblasts are the first cells recruited to the wound bed where they repopulate the ECM, while recruitment of papillary fibroblasts is associated with re-epithelialization and hair follicle generation ^11^.

These subpopulations also display heterogeneous responses in fibrosis ^13–16^. Nonetheless, the different contributions of various fibroblast subpopulations in response to pro-fibrotic stimuli and the molecular mechanisms to explain their differential activities remain largely unknown. To begin to uncover these molecular mechanism, we utilized a Snail transgenic (Snail Tg) mouse model of skin fibrosis that mimics the overexpression of this transcription factor found in the epidermis of scleroderma (SSc) patients ^17,18^. Ectopic expression of Snail is sufficient to induce phenotypes that recapitulates many of the diagnostic features of systemic scleroderma ^18^. It is also interesting to note that scleroderma patients also have higher incidence of developing cancers ^19^, suggesting that insights into this fibrotic disease would likewise shed light on the factors driving tumorigenesis. In line with this, the Snail Tg mice have also been shown to prime the skin towards the development of cutaneous squamous cell carcinoma ^20,21^. The mesenchymal compartment of a fibrotic tissue has remarkable similarities to the stroma surrounding solid tumours ^22,23^. The fibrosis associated with tumour stroma is driven by cancer-associated fibroblasts (CAFs) ^24^. Much like their counterparts in normal tissue, CAFs are heterogeneous in nature ^23^. One subpopulation of CAFs, known as myofibroblastic CAFs (myCAFs) expresses higher levels of α-Smooth Muscle Actin (αSMA), collagens and other genes associated with myofibroblast functions ^25^. Another major subpopulation is the inflammatory CAFs (iCAFs) which majorly expresses inflammatory cytokines ^25^. Collectively these different CAFs promote tumour associated inflammation, ECM remodelling of stroma, regulate cancer cell metabolism, survival and maintenance of cancer stem cells, promote metastasis, and aid in chemoresistance ^22,23^. Despite their important contributions to tumorigenesis, the etiology of these different CAFs remains unanswered. We have previously shown that Snail Tg keratinocytes secrete a factor Mindin in absence of which fibrogenesis and skin inflammation is abrogated ^18^. This presented us with a unique opportunity of further understanding the role of Mindin in elucidating the regulation of the functional heterogeneity of fibroblasts in both tissue fibrosis and tumor stroma.

## RESULTS

### Snail Transgenic (Snail Tg) mouse has perturbed proportions and localization of fibroblasts subpopulations

We have previously shown that αSMA, a marker for activated fibroblasts, is upregulated in the Snail Tg dermis as early as postnatal day 9 ^26^. The dermal fibroblasts can be spatially subdivided into two major subpopulations, papillary fibroblasts in the upper dermis and reticular and hypodermal fibroblasts in the lower dermis ^12,27^. These subpopulations can be isolated based on surface markers, CD26 for papillary and Sca1 for lower reticular/hypodermal fibroblasts ^8,11,13^. The fibroblast heterogeneity parameters such as the proportion of specific fibroblast subtypes may be altered during repair and disease conditions ^28^. To determine which subpopulation of dermal fibroblasts are activated in the Snail Tg skin, we performed flow cytometry analysis using vimentin as an established marker of fibroblasts ^29,30^ co-stained with either Sca1/aSMA or CD26/aSMA. Both subpopulations (CD26+/Vim+ and Sca1+/Vim+) showed an increase in the number of activated cells in the Snail Tg background relative to its wild type (WT) control (Figure 1A). Interestingly, when we investigated the spatial organization of the cells, we found that Sca1+ fibroblasts are progressively recruited to the epidermal-dermal junction (Figure 1C-1D, S1D). Analysis of the probability distribution of Sca1+ fibroblasts as a function of distance from the epidermis, revealed that Sca1+ cells were distributed between 50-150 um from the epidermis in WT skin at all postnatal ages analysed (Figure 1C and Figure S1D). In the Snail Tg skin, the Sca1+ cells are first noticed to have a differential localization at P5 and the probability of finding these cells near the epidermis gradually increases with time. As we previously reported, dermal thickening in the Snail transgenic skin can be initially observed at P9 ^26^. This is consistent with the probability distribution wherein Sca1+ fibroblasts are also observed to be lower in the Snail Tg dermis at P9 relative to the WT skin. Interestingly the papillary localization CD26+ fibroblasts between the WT and Snail Tg skin remain largely invariant (Figure 1E-1F, S1E).

**Figure 1:**
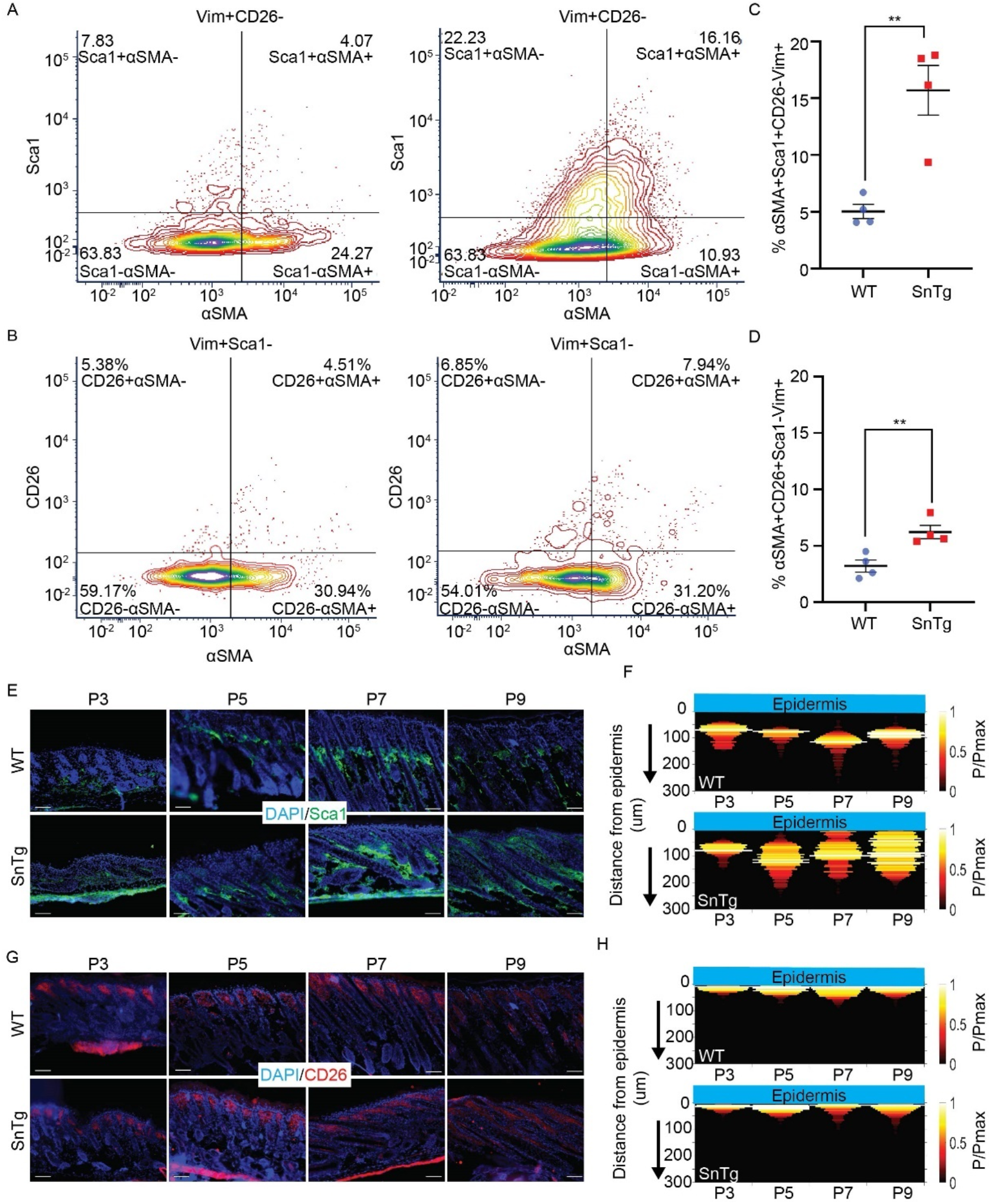
Sca1+ fibroblast localization is perturbed in the dermis of Snail transgenic (SnTg) mice. Representative contour plot showing quadrants for (A) αSMA±Sca1±CD26-Vim+ cells and (B) αSMA±CD26±Sca1-Vim+ cells, in postnatal day 9 (P9), WT (left panel) and SnTg (right panel). Individual value plot with mean±SEM of (C) % αSMA+Sca1+CD26-Vim+ and (D) % αSMA+CD26+Sca1-Vim+ cells (n=4; p-values were calculated by Welch’s t-test, ** P < 0.01). (E) Sca1+ fibroblasts (green) and nuclear staining with DAPI (blue) in WT and SnTg skin sections in postnatal day 3 (P3), P5, P7 P9 pups. (F) Heatmap showing the probability of Sca1+ cells at a given distance below the epidermis in WT (top panel) and SnTg (Bottom panel) mice at P3, P5, P7, and P9. (G) CD26+ fibroblasts (green) and nuclear staining with DAPI (blue) in WT and SnTg skin sections from P3, P5, P7, and P9 pups. (H) Heatmap showing the probability of CD26+ cells at a given distance below the epidermis in WT (top panel) and SnTg (Bottom panel) at P3, P5, P7, and P9.

### Mindin induces migration of Sca1+ fibroblasts

We recently observed that the matricellular protein Mindin (Spondin-2) is secreted from Snail Tg keratinocytes ^18,21^ and is required for the expression of inflammatory cytokines in dermal fibroblasts and fibrosis in the transgenic mouse ^18^. To test if Mindin is also required for the relocalisation of Sca1+ fibroblast towards the epidermal-dermal junction in the Snail Tg skin we stained Sca1+ fibroblasts (Figure 2A) and quantified (Figure 2B, S2A) their localization in P9 WT, Snail Tg, and Snail Tg/Min KO (Mindin knockout) skin. The localisation of Sca1+ cells in Snail Tg Min KO was similar to WT skin, indicating that Mindin is required for the relocalization of the cells from the lower dermis towards the epidermal-dermal junction. To test if Mindin is sufficient to change Sca1+ fibroblast localisation, we tested if Mindin can function as a chemoattractant in vitro. For this purpose, we sorted and cultured CD26+ and Sca1+ fibroblasts and seeded them in a transwell chamber. Upon addition of Mindin to the lower chamber of the transwell, unsorted fibroblasts, and Sca1+ fibroblasts migrated to the bottom, but not CD26+ fibroblasts (Figure 2C). Indicating that the chemotactic response to Mindin is unique to Sca1+.

**Figure 2:**
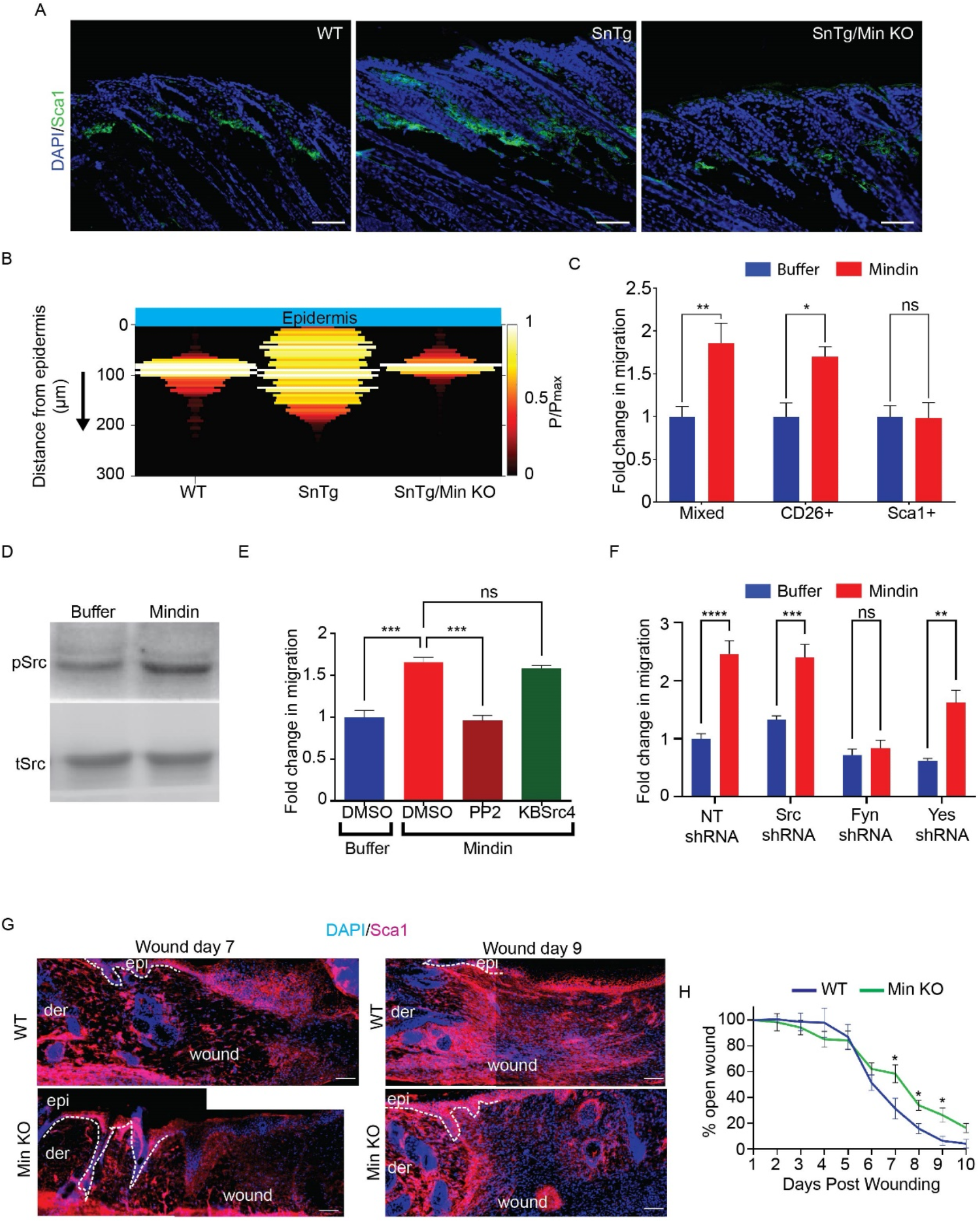
Mindin induces migration of Sca1+ fibroblasts via Fyn kinase. (A) IF staining for Sca1 in P9 WT, SnTg and SnTg/Mindin knockout (Min KO) skin (scale bar = 50 µm). (B) Heatmap showing probability of Sca1+ cells at a given distance below the epidermis in WT, SnTg and SnTg/Min KO skin. WT and SnTg are the same data as in Figure 1. (C) Transwell assay to measure migration of mixed, CD26+ or Sca1+ fibroblasts with either buffer or Mindin as a potential chemoattractant (n≥4). (D) Amount of phosphorylated Src (pSrc) and total Src proteins (tSrc) in fibroblasts treated with either buffer or Mindin for 15 mins. (E) Transwell assay with Sca1+ fibroblasts stimulated with buffer or Mindin in the presence of DMSO, PP2 (10 µM), or KbSrc4 (10 µM) (n=3). (F) Transwell assay with. Sca1+ fibroblasts transduced with either Non-targeting (NT), Src, Fyn or Yes shRNA with buffer or Mindin as a chemoattractant (n=3). (G) IF for Sca1, in WT and Min KO day 7 and day 9 skin wounds (The images were stitched manually; scale bar = 50 µm) (H) Percent wound closure in WT and Min KO mice w.r.t. wound size on day 1. (n = 3 mice, 2 wounds per mice). Data represents the mean ± SEM. p-values were calculated by Welch’s t-test (C and H), 1-way ANOVA followed by Tukey’s post hoc analysis (E), 2-way ANOVA followed by post hoc Šídák’s multiple comparisons test (F) (*P < 0.05, **P < 0.01, ***P < 0.001, ****P < 0.0001 and ns (P>0.05) is non-significant).

To achieve insights into the molecular mechanisms underlying the pro-migratory effect of Mindin on cells we performed GOTERM enrichment analysis on differentially upregulated genes (1715 upregulated genes with P<0.05, FC>1.5 folds) from RNAseq data of Mindin treated fibroblasts ^18^. The analysis revealed significant enrichment in biological processes associated with cell migration (41 genes), and positive regulation of cell migration (45 genes) (Figure S2B). We created a sublist of 81 upregulated genes and reperformed GSEA. We found enrichment in processes associated with the cytoskeletal organization, cell adhesion, ECM organization, along with the activation of integrin signalling and kinases involved in cell migration (Figure S2C). Mindin is a known integrin ligand ^31–33^ and we hypothesized that Mindin may activate the Src family of Kinases (SFK), given that the integrin-SFK axis has been previously reported to mediate fibroblast migration ^34–36^. To test whether Mindin exposure activates SFK in dermal fibroblasts we probed for phospho-Src (pSrc) levels via western blot. We observed that Mindin was able to activate Src family kinases within 15 minutes of treatment (Figure 2D and S2D). To assess whether the activation of Src family kinases by Mindin is necessary for migration of Sca1+ fibroblasts, we used two different SFK inhibitors - pp2 (pan SFK inhibitor) ^37,38^ and KbSrc4 (a preferential inhibitor of c-Src) ^39^. While addition of pp2 to the transwell chamber inhibited the migration of Sca1+ cells, this was not the case with KbSrc4 (Figure 2E). Given that KbSrc4 is more potent in inhibiting c-Src over other SFK members such as Fyn and Yes ^39^, this indicated that there might be a differential role of SFK members in mediating the effect of Mindin. Thus, to delineate this further, we generated shRNA based knockdowns of Src, Fyn and Yes kinases which showed 60-80% reduction in RNA expression of Src, Fyn and Yes, respectively (Figure S2E). In a transwell migration assay, Sca1+ cells transduced with non-targeting shRNA, Src shRNA and Yes shRNA migrated in response to Mindin, but cells transduced with Fyn shRNA did not (Figure 2F). This indicated a non-redundant essential role of Fyn in migration of Sca1+ fibroblasts downstream of Mindin.

Given the evidence for the role of Mindin in fibroblast migration, we investigated whether Mindin is also important for localisation/recruitment of Sca1+ fibroblast in a physiological context such as wound healing. We wounded WT mice and quantified the expression of Mindin RNA in the wound tissue from day 1 to day 10 post wounding. We observed that Mindin expression starts increasing from day 3 post wounding and peaks at day 7 and decreases thereafter (Figure S2F). We therefore stained the wound tissue for Sca1+ cells on day 7 and day 9 post wounding. Immunofluorescence staining of day 7 and day 9 post wounded skin revealed decreased numbers of Sca1+ cells localised in the wound bed in Min KO compared to WT mice (Figure 2G). In line with this, we also observed a significant difference in percentage of wound closure at post wounding day 7 - day 9 (Figure 2H).

### Mindin induces an inflammatory phenotype in Sca1+ fibroblasts

We have previously reported that there is an increase in inflammation in Snail Tg skin ^26,40^. Furthermore, we observed that Mindin deficiency in the Snail transgenic background resulted in a decrease in cutaneous inflammation ^18^. In addition, purified Mindin was sufficient to induce inflammatory cytokine expression in fibroblasts in vitro ^18^. We then analysed which fibroblast subpopulation was responsive to Mindin to induce an inflammatory response. Hence, we treated the sorted population with recombinant Mindin in vitro and quantified the expression of various cytokines which were observed to be differentially upregulated in the RNAseq analysis of Mindin treated fibroblasts such as Rantes (CCL-5), IL-6, CXCL-10, CXCL-5, CXCL-3. and IL17. In CD26+ fibroblasts Mindin did not cause significant upregulation of Rantes, CXCL-10, CXCL-5 or IL17, however, there was a small but significant increase in IL-6 expression (Figure 3A). On the other hand, Mindin treated Sca1+ cells exhibited a robust upregulation of all cytokines (Figure 3B). Furthermore, CXCL-3 which was undetected in CD26+ fibroblasts was expressed in Sca1+ fibroblasts and showed a robust increase in response to Mindin. This data indicates that Mindin is sufficient to strongly induce the expression of pro-inflammatory cytokines in Sca1+ fibroblasts.

**Figure 3:**
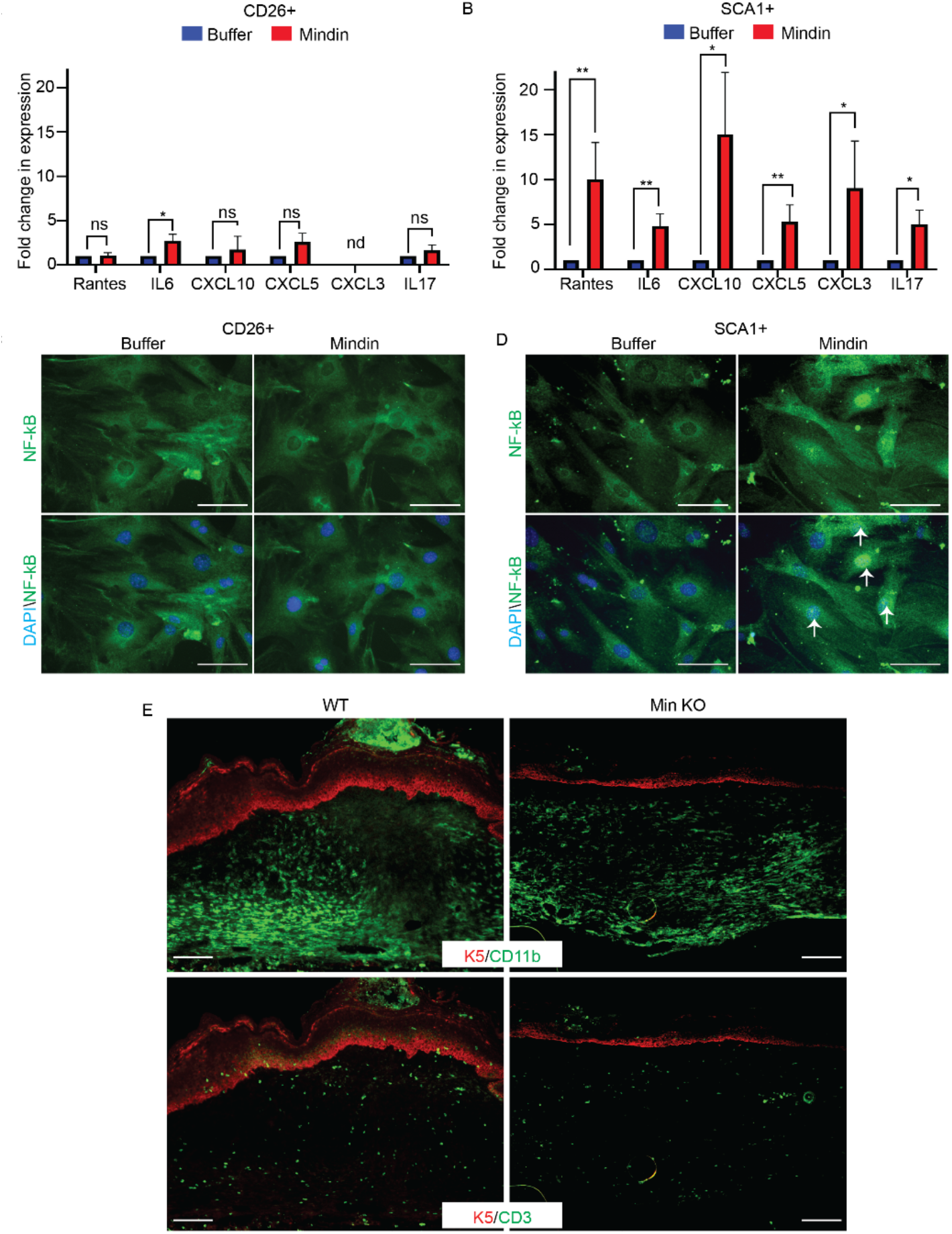
Mindin stimulates inflammatory cytokine production in Sca1+ fibroblasts. qPCR for expression of inflammatory cytokines from (A) CD26+ fibroblasts or (B) Sca1+ fibroblasts treated with either buffer or Mindin (n≥4). Data represents the mean ± SEM. p-values were calculated by Ratio paired t-test (A and B) (*P < 0.05, **P < 0.01, ***P < 0.001, ****P < 0.0001 and ns (P>0.05) is non-significant and nd=not detected). IF staining for NF-κB (green) and DAPI (blue) in CD26+ fibroblasts (C) or Sca1+ fibroblasts (D) treated for 1-hour with either buffer or Mindin. (E) IF staining for CD11b (macrophages; top panels) and CD3 (T-Cells; lower panels) in WT(left) and Min KO (right) skin sections post-wounded day 7 (scale bar = 50 μm).

We then focused on the mechanism by which extracellular Mindin can stimulate cytokine gene expression. Given that Mindin has been previously shown to activate the NF-kB pathway in renal cells (HK-2 cells) treated with TGF-β ^41^, we investigated if NF-kB signalling mediates the inflammatory response in fibroblast following induction by Mindin. In line with this we found the NF-kB pathway as one of the KEGG pathways enriched in the GSEA using upregulated genes from Mindin treated fibroblasts (Figure S3A). Moreover, we observed that one-hour treatment of a mixed population of dermal fibroblasts induced nuclear translocation of NF-kB in a subset of cells (Figure S3B). Together with other inflammatory pathways from the KEGG pathway analysis, we hypothesised that Mindin may stimulate inflammation in Sca1+ cells by activating the NF-kB pathway. To test this hypothesis, we isolated CD26+ cells and Sca1+ cells and treated them with recombinant Mindin and assayed for NF-kB subcellular localization. Even though CD26+ cells retained NF-kB in the cytoplasm with and without Mindin treatment (Figure 3C), the majority of Sca1+ cells demonstrated a nuclear translocation of NF-kB upon Mindin treatment (Figure 3D).

Given our observation that there is deficiency of Sca1+ cells in the wound bed at day 7 post wounding. We predicted that Mindin KO wounds will also have deficiency in recruiting immune cells into the wound bed. Consistent with this prediction, staining of day 7 post-wounding revealed a reduced number of T-cells in the wound beds of Mindin null animals, though macrophages were largely unaffected (Figure 3E).

### Mindin induces collagen contraction in CD26+ fibroblasts in a c-Src dependent manner

Though there was not an obvious perturbation in the localisation of CD26+ fibroblasts, a qualitative change in both a denser packing and a more uniform orientation of papillary fibroblasts was observed in the Snail Tg skin, which was lost in the absence of Mindin (Figure S4A). To test whether there is denser packing of the papillary fibroblasts, the distance between two neighbouring CD26+ cells was measured and plotted as a function of the distance from the epidermis (Figure 4A). In agreement with the qualitative observation, the quantification revealed a significant reduction in intracellular distance between neighbouring CD26+ cells in Snail Tg mice, closer to the epidermis. Nevertheless, in Snail Tg/Min KO mice the dense packaging of CD26+ cells, was attenuated and there was no significant difference between Snail Tg/Min KO and WT skin. Many factors can influence the density of cells in the dermis, one of which is the increased number of CD26+ cells in the Snail transgenic dermis. However, there was no difference in the number of papillary fibroblasts in the WT vs. Snail Tg skin (Figure S4B). Another possibility is that myofibroblast contraction may result in the compaction of the matrix resulting in an increased localised density of cells. Furthermore, fibroblasts in contracting gels become parallelly aligned and are closely packed together ^42^. Hence, we hypothesised that CD26+ fibroblasts may differentiate to contractile myofibroblasts in Snail Tg skin in a Mindin dependent fashion. To test this hypothesis, heterogeneous fibroblasts and FACS sorted Sca1+ and CD26+ fibroblasts were embedded in collagen gels and treated with Mindin. 72-hours post-treatment we measured the area of the treated gels and found that Mindin can promote collagen contraction in gels seeded with either mixed fibroblasts or CD26+ fibroblasts (Figure 4B). However, this was not the case for Sca1+ fibroblasts.

**Figure 4.**
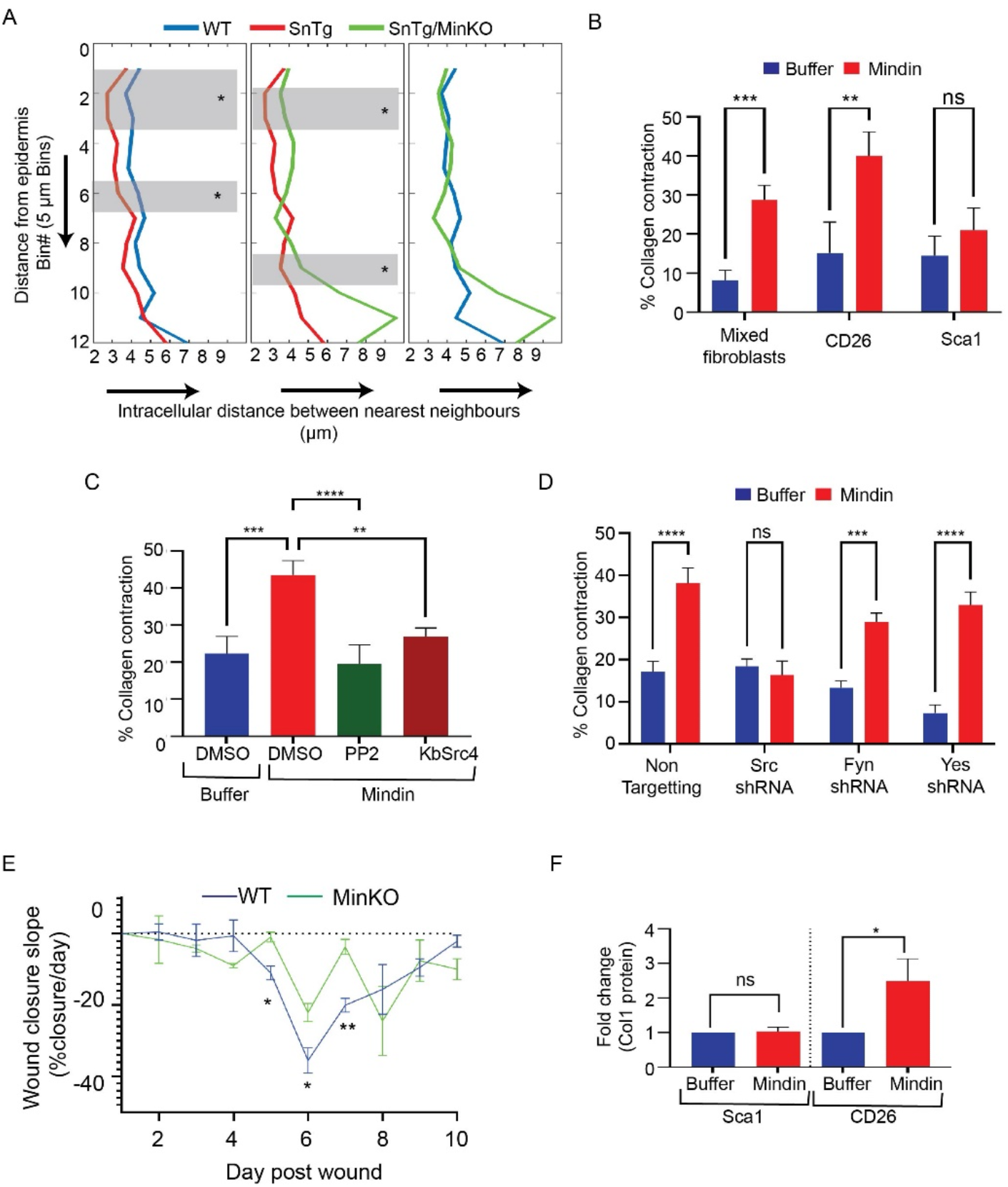
Mindin induces fibroblast contraction and collagen production in CD26+ fibroblasts. (A) Measurement of intracellular distance between two nearest CD26+ nuclei (x-axis) as a function of distance below the epidermis (y-axis, bin# below the epidermis, bin size = 5 µm) in WT, SnTg and SnTg Min KO skin. (n = 3. Number of CD26+ cells counted per skin section > 80. Region shaded in gray marks the bins where p<0.05) (B) Collagen contraction assay, showing percentage contraction of collagen gels seeded with either mixed, CD26+, or Sca1+ fibroblasts and treated with either buffer control or Mindin (n≥4). (C) Effect of SFK inhibition on Mindin induced collagen contraction. CD26+ fibroblasts were treated with either buffer control or Mindin along with either DMSO, PP2, or KbSrc4 (n≥3). (D) Effect of Non-targeting (NT), c-Src, Fyn, or Yes shRNA on collagen contraction with CD26+ fibroblasts, treated with either buffer control or Mindin (n≥3). (E) Measurement of slope of % wound closure/day in WT and Min KO mice. The slope was calculated as: % closure of given day - % closure on previous day (n = 3 mice, 2 wounds per mice). Quantification of Col1 in Mindin or buffer control treated CD26+ and Sca1+ fibroblasts, normalised to Lam (n=3). Data represents the mean ± SEM. p-values were calculated by Welch’s t-test (B, E), ratio-paired t-test (F), 1-way ANOVA followed by Tukey’s post hoc analysis (C), 2-way ANOVA followed by post hoc Šídák’s multiple comparisons test (D) (*P < 0.05, **P < 0.01, ***P < 0.001, ****P < 0.0001 and ns (P>0.05) is non-significant).

We then investigated the mechanism by which Mindin can induce contraction in CD26+ fibroblasts. The Src family of kinases has been shown to be involved in regulation of myofibroblast differentiation and contraction downstream of TGF-β1 (Hu et al., 2014). The integrin-Src signalling axis has been implicated in the regulation of RhoA-ROCK activation which can affect stress fibre formation and contractile activity via Myosin light chain phosphorylation ^43^. Since Mindin mediated migration of Sca1+ cells via Fyn Kinase, we tested if SFKs are also required for Mindin mediated contraction of CD26+ cells. Both the pan-SFK inhibitor PP2 as well as the c-Src specific inhibitor, KbSrc4, blocked Mindin mediated contraction (Figure 4C). To further access which member of the SFK was required for contraction of CD26+ cells, we knocked down Src, Fyn and Yes kinases by transduction of specific shRNAs. The transduced cells were embedded in collagen gels and treated with Mindin. While CD26+ fibroblasts transduced with either non-targeting, Fyn or Yes shRNA contracted the collagen gels in response to Mindin, reduction of c-Src inhibited the Mindin mediated contraction (Figure 4D). This indicated that specific Src family kinase members are differentially utilised for different aspects of fibroblast activation.

One of the physiological roles of fibroblast contraction is its contribution towards the contraction of the wound bed, which aids in wound closure. Since we observed a significant delay in wound healing in Mindin null animals (Figure 2H), we calculated the slope of each day in WT and Min KO wounds as a proxy for in vivo tissue contraction. Our results show that contraction in WT mice starts at D5 and continues till day 7 post-wounding. Interestingly, the magnitude of the contraction is lower in the Min KO animal (Figure 4E). However, unlike Sca1+ cells, we did not observe any deficiency in recruitment of CD26+ fibroblasts in the wound bed of Min KO animals (Figure S4C). This indicates that while Mindin is not required for migration and recruitment of CD26+ cells, it plays a role in contraction of the wounds.

Apart from contraction, another consequence of fibroblast activation is the increase in collagen production. Thus, to test if Mindin can affect collagen production we treated the sorted fibroblasts with Mindin and measured the levels of Collagen 1 (Col1a1 and Col1a2) expression via western blot (Figure 4F and S4D). Only CD26+ cells exhibited an increase in the amount of Col1 levels upon treatment with Mindin. Though Col1 is the most abundant collagen and contributor to tissue fibrosis there are other collagens which play important roles in tissue fibrosis - such as Col3, Col4, Col5, Col7 ^44,45^. The type of collagen upregulated may differ by the ligand and fibroblast subtype. Therefore, to test if Mindin differentially regulates expression of collagen sub-types in CD26+ and Sca1+ we measured the mRNA expression level of these collagens via qPCR. Col1a2, Col5a, and Col3a1 were significantly upregulated in Mindin treated CD26+ cells (Figure S4E, left panel). Interestingly, Col7a1, which was not significantly upregulated in CD26+ cells, was significantly increased in Mindin treated Sca1+ cells (Figure S4E, right panel). These results reveal how different fibroblast populations contribute in a complementary fashion to the overall increase in the amount of collagen proteins in the fibrotic tissue.

### Mindin promotes a CAF-associated self-renewal promoting capability in CD26+ fibroblasts

The mesenchymal compartment of a fibrotic tissue has remarkable similarities to the stroma surrounding solid tumours ^22,23^. We thus hypothesised that Mindin may have a role in the generation of cancer-associated fibroblasts, which plays many important roles in tumorigenesis ^23,25^. In support of the potential role of Mindin in cancers, Mindin is reportedly upregulated in many cancers and is proposed as a potential diagnostic and prognostic biomarker ^46–50^. Furthermore, gene set enrichment analysis using Mindin upregulated genes in fibroblasts, revealed significant enrichment of disease terms in the DisGeNET database, for inflammatory and fibrotic diseases and cancers (Figure S5A-S5B).

As shown in Figure 3A-B, Mindin differentially primes Sca1+ fibroblast to an inflammatory phenotype, which is consistent with iCAFs ^25^. In a similar vein, treatment of Mindin can endow CD26+ fibroblasts to become more contractile and secrete elevated collagen (Figure 4B and F), which is consistent with a myCAF phenotype ^25^. Furthermore, it has been shown that CD10, C5a, and GPR77 are signatures of a subset of CAFs in breast and lung cancer patients ^51^. Interestingly, only CD26+ fibroblasts increased expression levels of GPR77 and C5a upon Mindin treatment, though there was no effect on CD10 expression (Figure 5 A-B). C5a binds to its receptor GPR77 that activates NF-kB in a positive feedback loop to further increase expression GPR77 ^51^. In line with this, we found that prolonged treatment of CD26+ fibroblasts with Mindin can activate NF-kB signalling in these cells (Figure S5C).

**Figure 5:**
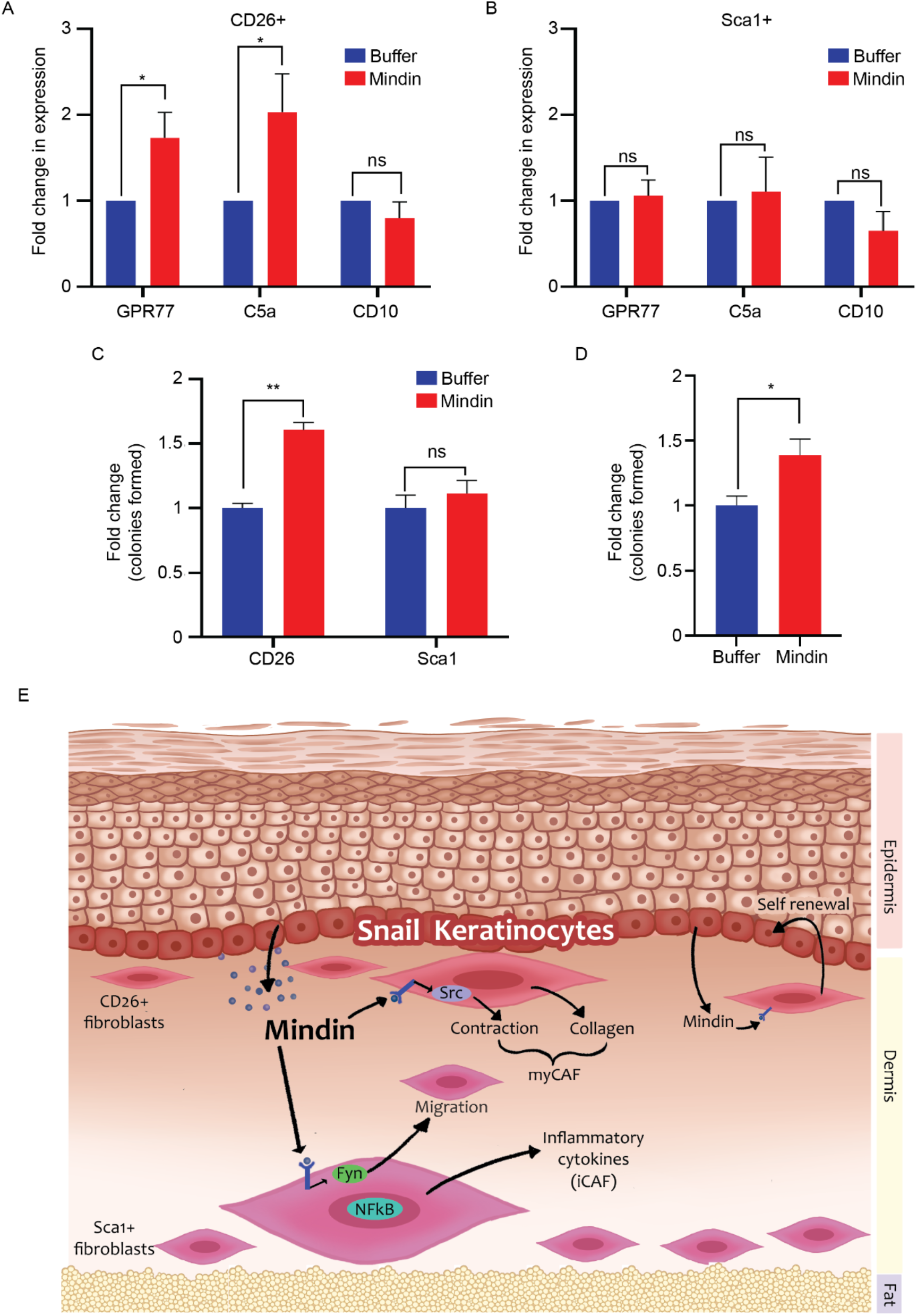
Mindin promotes CD26+ fibroblasts to adopt a CAF phenotype. Fold change in GPR77, C5a and CD10 expression in (A) CD26+ (n=6) and (B) Sca1+ (n=4) fibroblasts treated with either buffer of Mindin. (C) Colony formation assay for primary mouse keratinocytes co-cultured with CD26+ or Sca1+ fibroblasts pre-treated with either buffer of Mindin for 24-hours (n=3). (D) Colony formation assay for primary mouse keratinocytes cultured with conditioned media (CM) collected from CD26+, fibroblasts treated with either buffer of Mindin(n=4). Data represents the mean ± SEM. p-values were calculated by ratio paired t-test (A-C), Welch’s t-test (D-F), (*P < 0.05, **P < 0.01, ns (P>0.05) is non-significant). (E) Model of differential effects of Mindin on distinct subpopulations of dermal fibroblasts.

These CAFs are associated with maintenance of cancer stem cells (CSCs) and chemoresistance and therefore poor prognosis ^51^. We then tested whether Mindin-treated fibroblasts are functionally equivalent to CAFs, and in particular, able to promote self-renewal of epithelial progenitor/stem cells. For this purpose, we co-cultured primary mouse epidermal proliferating keratinocytes (mKTs) with CD26+ or Sca1+ fibroblasts pre-treated with either buffer or Mindin, and measured self-renewal in a colony formation assay. Only Mindin-treated CD26+ fibroblasts were able to increase colony formation of mKTs (Figure 5C). Furthermore, culturing mKTs in conditioned media (CM) collected from Mindin treated CD26+ fibroblasts was sufficient to increase colony formation (Figure 5D), indicating a role of molecular crosstalk between CD26+ dermal fibroblasts and keratinocytes by soluble factors.

## DISCUSSION

Our data reveals Mindin as a novel modulator of heterogeneous dermal fibroblasts that drive cutaneous fibrogenesis (Figure 5E). We have shown that Mindin is secreted by Snail Tg keratinocytes ^21^ and elicits unique functional responses in resident subpopulations of fibroblasts in the skin. Mindin mediates migration of Sca1+ fibroblasts via Fyn kinase and increases inflammatory cytokine production in these cells. On the other hand, Mindin induces a more contractile phenotype in CD26+ fibroblasts, and increases collagen 1 production in these cells. However, Mindin utilizes c-Src to mediate this effect in papillary fibroblasts. Interestingly, the effect of Mindin on Sca1+ and CD26+ fibroblasts endows them with features of iCAFs and myCAFs, respectively. In addition, Mindin-treated CD26+ fibroblasts are capable of promoting self-renewal of epithelial cells akin to the role of CAFs in maintaining cancer stem cells. This data is consistent with the notion that CAFs can be derived from resident fibroblasts within the issue ^52,53^.

We have found that the spatial organization of Sca1+ cells is perturbed in the Snail transgenic background and have attributed Mindin-mediated migration as an underlying cause for this. Besides cellular migration, another possible explanation is the possibility of CD26+ cells transdifferentiating into Sca1+ fibroblasts. One prediction of a transdifferentiating CD26+ to Sca1+ fibroblast would be the detection of a transitional state of the cell where they would be double-positive for both markers. However, both wild type and Snail transgenic skin contained only ∼2% of fibroblasts were double-positive for CD26+ and Sca1+ (Figure S6A and B), which cannot explain the increased number of Sca1+ cells at the dermal-epidermal junction in the transgenic animal. In addition to this, treatment with Mindin did not induce expression of Sca1 in CD26+ fibroblasts or vice versa (Figure S6C). A functional consequence of relocating the Sca1+ fibroblasts from the lower to the upper dermis is the deprivation of a potential source of adipose tissue that lie within the lower dermis. Sca1+ fibroblasts have been reported to express pre-adipocytes markers and contribute to the maintenance of adipose tissue homeostasis ^8,11,13^. Consistent with this, in the adult Snail transgenic skin there is a near total absence of dermal white adipose tissue that is replaced by collagen ^18^.

One notable observation was the increase in the number of Sca1+ fibroblasts in the Snail transgenic skin (Figure S6D) without an increase in proliferation (Figure S6E). As noted earlier, under homeostatic conditions, Sca1+ fibroblasts differentiate to maintain adipose tissue homeostasis ^11^. However, β-catenin stabilization in Sca1+ fibroblasts, has been shown to inhibit their differentiation into adipocytes thereby promoting fibrosis ^13^. Our results indicate that Mindin treatment activates β-catenin signalling in Sca1+ fibroblasts in-vitro, as measured by an increase in the β-catenin target gene Axin-2 (Figure S6F). This suggests that the increase in Sca1+ cells in the Snail transgenic skin is at least partly due to an accumulation of cells that do not otherwise differentiate into adipocytes. However, it cannot be ruled out that other fibroblast subpopulations such as Dlk+Sca1-reticular fibroblasts, which have been shown to contribute to all compartments of dermis ^11^, does not contribute to Sca1+ population in Snail Tg mice. Similarly, other multipotent sub-populations in the skin such as pericytes, invading fibrocytes, and mesenchymal stem cells (MSCs) may as well contribute to expansion of this compartment.

It has been shown that lower dermal fibroblasts are the major source of fibrous collagen during homeostasis and physiological repair ^12,27^. Nevertheless, we observed that it was the papillary fibroblasts which increased the levels of Col1 in response to Mindin. This suggests that in fibrotic scenarios, the CD26+ cells can be induced to provide a significant amount of ECM proteins. This is consistent with the report by Rinkevich et al.,2015 ^14^ that En1+ derived CD26+ fibroblasts constitute the major population with fibrogenic potential and CD26 inhibitor inhibits scarring. In addition to stimulating a local contracture, the ordered packaging of CD26+ papillary fibroblasts is also consistent with alignment of myCAFs observed immediately adjacent to the carcinoma ^54^. This packaging can arise from the contractile forces exerted on the matrix by myCAFs and remodelling of the local ECM ^54^. In addition to showing the features consistent with myCAFs, the papillary fibroblasts are therefore also in the histological location consistent with origins of myCAFs.

Mindin endows CD26+ cells with the ability to promote expansion/growth of the progenitor pool of epidermal keratinocytes akin to CAFs that promote self-renewal of CSCs 51. However, Mindin only upregulated 2 of the 3 molecular markers observed in CAFs - GPR77 and C5a, but not CD10. However as suggested by Su et al., 2018 51 CD10 is not essential for the downstream function of CAFs in promoting chemoresistance and self-renewal in these cells. NF-kB signaling has been shown to be important in mediating CAF activity, and it is interesting to note that while 1-hour treatment of Mindin in Sca1+ cells to induce nuclear localisation of NF-kB, but it took 24-hours in CD26+ fibroblasts. This suggests that NF-kB translocation may not be under direct regulation of Mindin in CD26+ cells but may depend on activation of other signalling pathways. Su et al., have shown that C5a and GPR77 interaction causes prolonged NF-kB activation in CAFs by acetylation of p65. This suggests that NF-kB activation in Sca1+ cells is via the canonical IKK pathway, whereas the activation in CD26+ cells is IKK-independent and thus has different kinetics.

Altogether, these findings provide new insights into the heterogeneous regulation of fibrosis-associated and cancer-associated fibroblasts and a number of interesting questions remain to be resolved. For instance, differential response of sub-population of fibroblasts corresponding to different members of SFKs might hint at differential expression receptors in these cells. These difference may be at the level of surface expression of distinct integrin pairs (which have been shown to be receptors for Mindin on macrophages, T-cells and colorectal cancer ^31,32,55^) and/or different cytoplasmic machinery downstream receptors in these subpopulations. Furthermore, there are many more sub-populations of fibroblasts that can contribute to fibrotic tissue which were not addressed in this study. These include Dlk-Sca1-reticular fibroblasts, pericytes, fibrocytes, and MSCs. Moreover, further refinement of these Sca1+ and CD26+ subpopulations and their responses to different pro-fibrotic stimuli may shed light into pathophysiology of fibrosis and the tumour stroma. For example myCAFs have been reported to be a double-edged sword and can function as both tumour promoting ^25^ or tumour-restraining ^56^. Thus it would be interesting to determine if refinement of CD26+ subpopulation can explain the two opposing phenotypes or if the same subpopulation can switch the phenotype in response to different microenvironmental compositions.

## METHODS

### Study approval

Animal work conducted at the NCBS/inStem Animal Care and Resource Centre was approved by the inStem Institutional Animal Ethics Committee following the norms specified by the Committee for the Purpose of Control and Supervision of Experiments on Animals (Government of India). All experimental work was approved by the inStem Institutional Biosafety Committee (IBSC).

### Animal studies

C57Bl6 (WT) mice were obtained from The Jackson Laboratory (Bar Harbor, Maine), Mindin KO mice were obtained from You-Wen He (Duke University), and the Snail Tg mouse was engineered as described earlier ^17^. The K14-Snail Tg/Mindin KO mouse was developed by breeding the K14-Snail Tg and Mindin KO mice.

### Cell culture

Primary newborn dermal fibroblasts (NBDF; Mixed fibroblasts) isolation was performed as described earlier ^26^ from C57Bl6 postnatal day 2 (P2)-day 3 (P3) pups. Fibroblasts were cultured in DMEM high glucose media with 10% FBS. For CD26+ and Sca1+ fibroblasts dermis was separated from the skin of P2 or P3 mice in 1 mg/ml of dispase for 1 h at 37 °C and digested in 2.5 mg/ml of collagenase IV (Gibco) for 1 h at 37 °C. Dermal cell suspension from 4-5 P2/P3 pups were pooled and stained with Sca1 (R&D FAB1226N; 1:30 μl) and CD26 (FAB9541P; 1:10 μl) for 30 minutes on ice. CD26+Sca1-(CD26+) and Sca1+CD26-(Sca1+) cells were sorted using FACS. The sorted cells (∼1 million) were seeded in a 3.5 cm dish and were cultured in the same manner as NBDF. All cells were maintained, and experiments were performed until passage # 5. Mouse epidermal keratinocytes were harvested from P2/P3 epidermis and were cultured in low calcium E-media to maintain undifferentiated proliferating state as previously described ^57^.

### Mindin purification and treatment

Histidine-tagged Mindin was purified from conditioned media collected from CHO-Mindin cells using Ni-NTA beads (Thermo Fisher Scientific) according to the manufacturer’s protocol. Varying concentrations of Imidazole (10-500mM) in 10mM tris, 300mM NaCl pH=8 was used for washing and elution. The purified Mindin was dialysed in buffer (10mM tris. 20mM NaCl, pH = 8) and filtered with 0.2 μm syringe filter. The purified recombinant Mindin was used at concentration of 80-200 ng/ml in all treatments. All treatments were done in serum-free conditions, unless otherwise stated.

### Fluorescence-activated cell analysis

P9 pups were sacrificed, and all hair was removed using Veet hair removal cream. Dermal cell suspension was prepared by separating dermis from epidermis in 1 mg/ml Dispase overnight at 4°C. Dermis was digested in 2.5 mg/ml of collagenase IV (Gibco) for 1 h at 37 °C. Dermal cell suspension was stained with Sca1 (R&D FAB1226N; 1:30 μl) and CD26 (FAB9541P; 1:10 μl) for 30 minutes on ice, fixed and permeabilized. They were then stained with primary anti-Vimentin (Abcam ab24525; 1:200) and anti-αSMA (Sigma A2547; 1:400) for 30 minutes. Ki67(Abcam AB16667) was used at 1:300. Secondary antibodies (Jackson ImmunoResearch) were used at 1:100. The cells were recorded using BD FACS ARIA Fusion and data was analysed on BD FACS Diva and FCS Express 7.0.

### Lentiviral transductions

All lentiviral production and transductions were done in BSL2 in accordance with IBSC guidelines. The method for lentiviral particle generation and the shRNA-pZip-mEf1a plasmids for c-Src, Fyn and Yes shRNA (procured from transomics), have been described earlier ^21^. The titre of virus that yielded greater than 50% GFP+ cells was used to transduce sorted Sca1+ or CD26+ fibroblasts after first passage, at 70-80% confluency. 72-hours post infection, the virus containing media was removed and fresh media with 1 μg/ml puromycin was added to enrich for transduced cells. Plates showing greater than 90% GFP+ cells were used for further experiments.

### Transwell migration assay

NBDF, CD26+, Sca1+, or shRNA transduced fibroblasts (50,000-100,000) were added to the upper chambers of 8-μm Transwell chambers in 10% FBS-containing DMEM and the same media was added to the bottom and incubated at 37 deg C for 2-4 h for cell attachment. Post incubation transwell chambers were washed with PBS and fresh serum-free media was added to the top. Transwell were then placed in fresh wells containing serum-free media with either control buffer or Mindin. For inhibitor assays serum-free media with either DMSO, PP2 (10 µM) (Millipore 529573) or KB SRC 4 (10 µM) (R&D 1008345) was added to the top 5 mins before placing the wells in Mindin containing serum-free media. 24h later the experiment was terminated and transwells were stained with crystal violet (CV) solution (0.5% CV, 10% ethanol, 4% PFA in PBS) and imaged after cleaning the top chamber.

Fold change in migration was calculated as:

(Number of cells/field migrated in the treated well)/(Number of cells/field in a migrated in the control well)

### Collagen contraction assay

Rat tail collagen (MilliporeSigma; 08-115) was dissolved in 0.1% acetic acid to make the concentration to 3 mg/ml stock. Cell suspension of NBDF, Sca1+, CD26+, or shRNA transduced fibroblasts was made in 0.5% serum media with 150,000 cells/ml. The cells and gels were made by mixing collagen stock and gels in 1:2 ratio (final concentration - 1mg/ml collagen, 100,000 cells/ml) along with simultaneous addition of an appropriate amount of 1M NaOH (predetermined via titration, ∼8-10µl/ml). Cell-free gels were made by mixing collagen stock and media without cells at 1:2 ratio. The mixture was immediately added to either 24 well (500 µl/well), 48 well (250 µl/well) or 96 (100 µl/well) well dish and incubated at 37 deg C for 1 h for solidification. Post solidification serum-free media (same volume as the gel) along with either buffer or Mindin was added to well. For inhibitor assays PP2 (10 µM) or KB SRC 4 (10 µM) was supplemented along with buffer or Mindin. The gels were detached from the walls using a pipette tip and kept at 37 deg C. 72 h later the experiment was terminated, and gels were stained with CV solution for contrast enhancement and images were acquired for quantification. The percentage contraction was calculated as: percent contraction, C = (1 – At/Anc)*100, where At is area of collagen gel containing cells and with treatments and Anc is the area of collagen gel with no cells added, as measured 72-hours post the detachment step.

### Western blot

Cells were serum-starved overnight before treatment and were treated with either buffer or Mindin in serum-free medium for 15 minutes. Cells were lysed in RIPA buffer and were loaded after addition of Laemmli buffer. The membranes were probed for phosphorylated Src (pSrc; CST 2101) and then with total Src (tSrc; CST2123) after stripping. For assessment of collagen 1, serum-starved cells are treated with either buffer or Mindin for 48-hours. Cells were lysed in RIPA and the lysates were loaded in the gels along with Laemmli buffer. The membranes were probed with anti-collagen antibodies (Abcam ab21286) and Lamin B (Abcam AB16048). The HRP-labelled secondary antibodies (Jackson ImmunoResearch) were used at 1:3,000 dilution. Blots were developed on an ImageQuant LAS4000, and bands were quantified using Fiji software (ImageJ, NIH).

### Cell localisation analysis

The algorithm for calculating the spatial probability distribution is described on GitHub (https://github.com/skinlab-sunnyk/celllocalization). Briefly, the images were rotated to make epidermis parallel to the horizontal axis. The X and Y coordinates (in pixel) were derived by marking the nuclei in the epidermal compartment, and nuclei positive for either Sca1 or CD26 in the dermal compartment, using Fiji Image J multipoint tool ^58^. These coordinates were entered into the algorithm which calculated the distance of a given cell in the dermis from its nearest cell in the epidermis and assigned a bin number based on this distance. Empirical probability of finding a cell as a function of distance from epidermis as:

P(cell type, bin number) = number of cells in a bin/total number of cells counted.

Bin size = 5 µm was used based on average size of nucleus in dermis. Welch’s t-test was used to compare means of corresponding bins between WT and Snail Tg, Snail Tg and Snail Tg/Min KO, or WT and Snail Tg/Min KO mice. To visualise how the likelihood of finding a cell relative to expected niche, spatial probability P was scaled with its maximum value (P/Pmax). P/Pmax was used to generate the heatmap, where the width and intensity were maximum when P/Pmax = 1.

### Nearest neighbour analysis

The algorithm for calculating the spatial nearest neighbour distances w.r.t. distance from epidermis is shared on GitHub (https://github.com/skinlab-sunnyk/nnanalysis). Briefly, the images were rotated to make epidermis parallel to the horizontal axis. The X and Y coordinates (in pixel) were derived by marking the nuclei in the epidermal compartment, and nuclei positive for CD26 in the dermal compartment using Fiji Image J multipoint tool ^58^. These coordinates were entered into the algorithm which calculated the distance of a given cell in the dermis from its nearest cell in the epidermis and assigned a bin number based on this distance. The distance matrix was created with the distance of each CD26+ cell with every other CD26+ cell in a given section. The algorithm then extracts the distance to the nearest neighbour of each cell. The average distance to nearest neighbour in a given bin for a given section is calculated as, 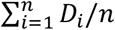, where Di is distance to nearest neighbour of i^th^ cell in a given bin and n is number of cells in that bin. Bin size = 5 µm was used based on average size of nucleus in dermis. Welch’s t-test was used to compare corresponding between WT and Snail Tg, Snail Tg and Snail Tg/Min KO, or WT and Snail Tg/Min KO mice.

### Colony formation

For co-culture assays, CD26+ and Sca1+ fibroblasts were seeded in a 3.5 mm dish at low density (∼10,000 cells). 24-hours post-seeding cells were serum starved overnight and then treated with either buffer or Mindin. 24-hours post-treatment the media was removed, cells were washed with PBS, and 1000 undifferentiated keratinocytes were added per dish along with low calcium E-media. 7 days post adding keratinocytes the cells were fixed and stained with crystal violet for counting colonies. For conditioned media experiments, CD26+ fibroblasts were seeded in a 12 well dish. At ∼80%-90% confluency cells were serum-starved and then treated with either buffer or Mindin for 24-hours. 24-hours post-treatment the media was removed, and fresh low Calcium E-media was added. 48-hours later the conditioned media was collected. Keratinocytes were seeded in a 24 well format (100 cells/well). 24-hours post-seeding the media was changed with 1:1 mixture of fresh low calcium E-media and conditioned media. 7 days post adding conditioned medium the cells were fixed and stained with crystal violet for counting colonies.

The fold change was calculated as: colonies counted in treated/average colonies counted in control

### Wound healing

Two excisional wounds (separated by 1-1.5) cm were created on the mid-dorsal region skin of the anaesthetised 3-4 month old mice. Pictures were taken from day 1 - day 10 post wounding. Percent wound closure was calculated as C = (1 - Wn/W1)*100, where C is percent wound closure, Wn is wound area on day n, and W1 is wound area on day 1. Slope or rate of wound closure was determined as % closure/day, R = Cn-Cn-1, where Cn is % closure on a given day n and Cn-1 is % closure on the previous day. Mice were sacrificed, and wounds were harvested on day 0 (unwounded), day 3, day 5, day 7, and day 9 and day 10 (wound closed), and tissue was stored in RNA later (for gene expression analysis) or embedded in OCT for immunofluorescence.

### NF-kB nuclear localisation

Cells were seeded in a 96 well dish (10000 cells/well) and were serum-starved overnight 24-hours later, followed by treatment with either buffer or Mindin. Post-treatment cells were fixed with 4% PFA, permeabilized and stained with NF-kB (Santa Cruz SC372) antibody at 1:200 dilution. Secondary antibody (Jackson ImmunoResearch) was used at 1:200. DAPI stain was used to mark nuclei. Images were captured with an Olympus IX73 microscope.

### Immunofluorescence of skin sections

Skin tissues were fixed and sectioned as previously reported ^40^, and probed with the following antibodies diluted 1:200: K5 (generated in-house); Sca1 (R&D AF1226); CD26(AF954); CD11b (Abcam ab8878); CD3 (e-biosciences 14-0041-85). Secondary antibodies (Jackson ImmunoResearch) were used at 1:200. DAPI stain was used to mark nuclei. Images were captured with an Olympus IX73 microscope.

### Gene set enrichment analysis (GSEA)

Gene list was created using the upregulated genes with adjusted p-value(q)<0.05 and fold change > 1.5 from RNAseq dataset of Mindin treated human dermal fibroblasts described earlier ^18^. Database for Annotation, Visualisation and Integrated Discovery (DAVID) was used for GSEA ^59–61^. GOTERM_BP_Direct ^62,63^was used to visualise enriched biological processes, KEGG pathways ^64^ for identifying associated signalling pathways and DisGeNET database for identifying the enrichment in associated diseases ^65^.

### Gene Expression

Total RNA was extracted from cells treated with either buffer or Mindin for 16-hours using TRIzol Reagent (TaKaRa, Thermo Fisher Scientific) and cDNA was synthesised using Superscript III (Thermo Fisher Scientific). The quantitative PCR (qPCR) was performed using Power SYBR Mix (Life Technologies, Thermo Fisher Scientific) in a Bio-Rad CFX384 machine. GAPDH or Actin expression was used as a reference for normalisation. The primer sequences used are listed in Supplemental Table 1.

### Statistics

Welch’s t-test was used for comparison of two groups. Ratio-paired t-test was used for comparison of fold changes in paired data. One-way ANOVA followed by Tukey’s post hoc analysis was used for comparing three or more groups. 2-way ANOVA followed by post hoc Šídák’s multiple comparisons test was used to compare 3 or more groups over multiple conditions. GraphPad Prism 6 (GraphPad Software) was used for all statistical analyses.

Data is represented as the mean ± SEM. p-values of less than 0.05 were considered significant.

## ACKNOWLEDGEMENTS

The authors would like to thank members of Jamora laboratory for their critical review of the work and insightful discussions, and Achyuth Acharya for their technical assistance, and Ritoparna Hazra for designing the graphical model. This work was supported by core funds from inStem, and grants from the Department of Biotechnology (DBT) of the Government of India (BT/PR8738/AGR/36/770/2013) and (BT/PR32539/BRB/10/1814/2019); the NIH/NIAMS (5R01AR053185-03); and the American Cancer Society (15457-RSG-08-164-01-DDC) to CJ. SK and KB were partially supported by the National Centre of Biological Sciences. IR was supported by the Indian Council of Medical Research (Senior Research Fellowship); Animal studies were partially supported by the National Mouse Research Resource (NaMoR) grant BT/PR5981/MED/31/181/2012;2013-2016;2018 and 102/IFD/SAN/5003/2017-2018 from the DBT. We thank the staff of the BLiSC Animal Care and Resource Centre and the BLiSC Central Imaging and Flow Cytometry Facility for technical assistance

## AUTHOR CONTRIBUTIONS

SK and CJ conceptualised and designed experiments, evaluated and interpreted data, and wrote the manuscript. SK, IR, KB, RFZ, GK, RZ, BD performed experiments. PK, SK performed bioinformatics analysis. AG provided resources and guidance for experimental design and analysis. CJ provided guidance and reviewed the manuscript.

## REFERENCES

1. Wynn T. Cellular and molecular mechanisms of fibrosis. J Pathol. 2008;214(2):199–210. doi:10.1002/path.2277

2. Thannickal VJ, Zhou Y, Gaggar A, Duncan SR. Fibrosis: Ultimate and proximate causes. J Clin Invest. 2014;124(11):4673–4677. doi:10.1172/JCI74368

3. Kendall RT, Feghali-Bostwick CA. Fibroblasts in fibrosis: Novel roles and mediators. Front Pharmacol. 2014;5 MAY:123. doi:10.3389/FPHAR.2014.00123/BIBTEX

4. Thiruvoth FM, Mohapatra DP, Kumar D, Chittoria SRK, Nandhagopal V. Current concepts in the physiology of adult wound healing. Plast Aesthetic Res. 2015;2(5):250–256. doi:10.4103/2347-9264.158851

5. Bainbridge P. Wound healing and the role of fibroblasts. J Wound Care. 2013;22(8):407–412. doi:10.12968/JOWC.2013.22.8.407

6. Wynn TA. Common and unique mechanisms regulate fibrosis in various fibroproliferative diseases. J Clin Invest. 2007;117(3):524–529. doi:10.1172/JCI31487

7. Korosec A, Frech S, Gesslbauer B, et al. Lineage Identity and Location within the Dermis Determine the Function of Papillary and Reticular Fibroblasts in Human Skin. J Invest Dermatol. 2019;139(2):342–351. doi:10.1016/J.JID.2018.07.033

8. Philippeos C, Telerman SB, Oulès B, et al. Spatial and Single-Cell Transcriptional Profiling Identifies Functionally Distinct Human Dermal Fibroblast Subpopulations. J Invest Dermatol. 2018;138(4):811. doi:10.1016/J.JID.2018.01.016

9. Ghetti M, Topouzi H, Theocharidis G, et al. Subpopulations of dermal skin fibroblasts secrete distinct extracellular matrix: implications for using skin substitutes in the clinic. Br J Dermatol. 2018;179(2):381. doi:10.1111/BJD.16255

10. Tabib T, Morse C, Wang T, Chen W, Lafyatis R. SFRP2/DPP4 and FMO1/LSP1 define major fibroblast populations in human skin. J Invest Dermatol. 2018;138(4):802. doi:10.1016/J.JID.2017.09.045

11. Driskell RR, Lichtenberger BM, Hoste E, et al. Distinct fibroblast lineages determine dermal architecture in skin development and repair. Nature. 2013;504(7479):277–281. doi:10.1038/nature12783

12. Zou ML, Teng YY, Wu JJ, et al. Fibroblasts: Heterogeneous Cells With Potential in Regenerative Therapy for Scarless Wound Healing. Front Cell Dev Biol. 2021;9:1955. doi:10.3389/FCELL.2021.713605/BIBTEX

13. Mastrogiannaki M, Lichtenberger BM, Reimer A, Collins CA, Driskell RR, Watt FM. β-Catenin Stabilization in Skin Fibroblasts Causes Fibrotic Lesions by Preventing Adipocyte Differentiation of the Reticular Dermis. J Invest Dermatol. 2016;136(6):1130–1142. doi:10.1016/J.JID.2016.01.036

14. Rinkevich Y, Walmsley GG, Hu MS, et al. Skin fibrosis. Identification and isolation of a dermal lineage with intrinsic fibrogenic potential. Science. 2015;348(6232). doi:10.1126/SCIENCE.AAA2151

15. Suttho D, Mankhetkorn S, Binda D, Pazart L, Humbert P, Rolin G. 3D modeling of keloid scars in vitro by cell and tissue engineering. Arch Dermatological Res 2016 3091. 2016;309(1):55–62. doi:10.1007/S00403-016-1703-2

16. Nazari B, Rice LM, Stifano G, et al. Altered Dermal Fibroblasts in Systemic Sclerosis Display Podoplanin and CD90. Am J Pathol. 2016;186(10):2650. doi:10.1016/J.AJPATH.2016.06.020

17. Jamora C, Lee P, Kocieniewski P, et al. A Signaling Pathway Involving TGF-β2 and Snail in Hair Follicle Morphogenesis. PLOS Biol. 2004;3(1):e11. doi:10.1371/JOURNAL.PBIO.0030011

18. Rana I, Kataria S, Tan TL, et al. Mindin is essential for cutaneous fibrogenesis in a new mouse model of systemic sclerosis. bioRxiv. Published online January 28, 2022:2022.01.26.477822. doi:10.1101/2022.01.26.477822

19. Weeding E, Casciola-Rosen L, Shah AA. Cancer and scleroderma. Rheum Dis Clin North Am. 2020;46(3):551. doi:10.1016/J.RDC.2020.03.002

20. Du F, Nakamura Y, Tan T-L, et al. Expression of Snail in Epidermal Keratinocytes Promotes Cutaneous Inflammation and Hyperplasia Conducive to Tumor Formation. Cancer Res. 2010;70(24). Accessed April 15, 2017. http://cancerres.aacrjournals.org/content/70/24/10080.long

21. Badarinath K, Dam B, Kataria S, et al. Snail maintains the stem/progenitor state of skin epithelial cells and carcinomas through the autocrine effect of the matricellular protein Mindin. bioRxiv. Published online June 29, 2021:2021.06.26.450022. doi:10.1101/2021.06.26.450022

22. Piersma B, Hayward MK, Weaver VM. Fibrosis and cancer: A strained relationship. Biochim Biophys acta Rev cancer. 2020;1873(2):188356. doi:10.1016/J.BBCAN.2020.188356

23. Kalluri R. The biology and function of fibroblasts in cancer. Nat Rev Cancer. 2016;16(9):582–598. doi:10.1038/nrc.2016.73

24. Kalluri R, Zeisberg M. Fibroblasts in cancer. Nat Rev Cancer. 2006;6(5):392–401. doi:10.1038/nrc1877

25. Geng X, Chen H, Zhao L, et al. Cancer-Associated Fibroblast (CAF) Heterogeneity and Targeting Therapy of CAFs in Pancreatic Cancer. Front Cell Dev Biol. 2021;9. doi:10.3389/FCELL.2021.655152

26. Nakasaki M, Hwang Y, Xie Y, et al. The matrix protein Fibulin-5 is at the interface of tissue stiffness and inflammation in fibrosis. Nat Commun. 2015;6. doi:10.1038/ncomms9574

27. Driskell RR, Watt FM. Understanding fibroblast heterogeneity in the skin. Trends Cell Biol. 2015;25(2):92–99. doi:10.1016/j.tcb.2014.10.001

28. Shaw TJ, Rognoni E. Dissecting Fibroblast Heterogeneity in Health and Fibrotic Disease. Curr Rheumatol Rep. 2020;22(8). doi:10.1007/S11926-020-00903-W

29. Chang Y, Guo K, Li Q, Li C, Guo Z, Li H. Multiple Directional Differentiation Difference of Neonatal Rat Fibroblasts from Six Organs. Cell Physiol Biochem. 2016;39(1):157–171. doi:10.1159/000445613

30. Hsia LT, Ashley N, Ouaret D, Wang LM, Wilding J, Bodmer WF. Myofibroblasts are distinguished from activated skin fibroblasts by the expression of AOC3 and other associated markers. Proc Natl Acad Sci U S A. 2016;113(15):E2162–E2171. doi:10.1073/PNAS.1603534113/-/DCSUPPLEMENTAL

31. Li H, Oliver T, Jia W, He YW. Efficient dendritic cell priming of T lymphocytes depends on the extracellular matrix protein mindin. Embo J. 2006;25(17):4097–4107. doi:10.1038/sj.emboj.7601289

32. Li Y, Cao C, Jia W, et al. Structure of the F-spondin domain of mindin, an integrin ligand and pattern recognition molecule. EMBO J. 2009;28(3):286–297. doi:10.1038/emboj.2008.288

33. Zhang YL, Li Q, Yang XM, et al. Spon2 promotes m1-like macrophage recruitment and inhibits hepatocellular carcinoma metastasis by distinct integrin–rho gtpase–hippo pathways. Cancer Res. 2018;78(9):2305–2317. doi:10.1158/0008-5472.CAN-17-2867/653130/AM/SPON2-PROMOTES-M1-LIKE-MACROPHAGE-RECRUITMENT-AND

34. Cary LA, Chang JF, Guan JL. Stimulation of cell migration by overexpression of focal adhesion kinase and its association with Src and Fyn. J Cell Sci. 1996;109(7):1787–1794. doi:10.1242/JCS.109.7.1787

35. Klinghoffer RA, Sachsenmaier C, Cooper JA, Soriano P. Src family kinases are required for integrin but not PDGFR signal transduction. EMBO J. 1999;18(9):2459–2471. doi:10.1093/EMBOJ/18.9.2459

36. Zou L, Cao S, Kang N, Huebert RC, Shah VH. Fibronectin Induces Endothelial Cell Migration through β1 Integrin and Src-dependent Phosphorylation of Fibroblast Growth Factor Receptor-1 at Tyrosines 653/654 and 766. J Biol Chem. 2012;287(10):7190–7202. doi:10.1074/JBC.M111.304972

37. Hanke JH, Gardner JP, Dow RL, et al. Discovery of a Novel, Potent, and Src Family-selective Tyrosine Kinase Inhibitor: STUDY OF Lck-AND FynT-DEPENDENT T CELL ACTIVATION (*). J Biol Chem. 1996;271(2):695–701. doi:10.1074/JBC.271.2.695

38. Ma YC, Shi C, Zhang YN, et al. The Tyrosine Kinase c-Src Directly Mediates Growth Factor-Induced Notch-1 and Furin Interaction and Notch-1 Activation in Pancreatic Cancer Cells. PLoS One. 2012;7(3):e33414. doi:10.1371/JOURNAL.PONE.0033414

39. Brandvold KR, Steffey ME, Fox CC, Soellner MB. Development of a highly selective c-Src kinase inhibitor. ACS Chem Biol. 2012;7(8):1393–1398. doi:10.1021/CB300172E/SUPPL_FILE/CB300172E_SI_001.PDF

40. Pincha N, Hajam EY, Badarinath K, et al. PAI1 mediates fibroblast-mast cell interactions in skin fibrosis. J Clin Invest. 2018;128(5):1807–1819. doi:10.1172/JCI99088

41. Yang K, Li W, Bai T, et al. Mindin deficiency alleviates renal fibrosis through inhibiting NF-κB and TGF-β/Smad pathways. J Cell Mol Med. 2020;24(10):5740–5750. doi:10.1111/jcmm.15236

42. Nakagawa S, Pawelek P, Grinnell F. Long-term culture of fibroblasts in contracted collagen gels: Effects on cell growth and biosynthetic activity. J Invest Dermatol. 1989;93(6):792–798. doi:10.1111/1523-1747.EP12284425

43. Zent J, Guo LW. Signaling Mechanisms of Myofibroblastic Activation: Outside-in and Inside-Out. Cell Physiol Biochem. 2018;49(3):848–868. doi:10.1159/000493217

44. Karsdal MA, Nielsen SH, Leeming DJ, et al. The good and the bad collagens of fibrosis – Their role in signaling and organ function. Adv Drug Deliv Rev. 2017;121:43-56. doi:10.1016/J.ADDR.2017.07.014

45. Rudnicka L, Varga J, Christiano AM, Iozzo R V., Jimenez SA, Uitto J. Elevated expression of type VII collagen in the skin of patients with systemic sclerosis. Regulation by transforming growth factor-beta. J Clin Invest. 1994;93(4):1709–1715. doi:10.1172/JCI117154

46. Zhang Q, Wang X-Q, Wang J, et al. Upregulation of spondin-2 predicts poor survival of colorectal carcinoma patients. Oncotarget. 2015;6(17):15095–15110. doi:10.18632/oncotarget.3822

47. Yuan X, Bian T, Liu J, et al. Spondin2 is a new prognostic biomarker for lung adenocarcinoma. Oncotarget. 2017;8(35):59324. doi:10.18632/ONCOTARGET.19577

48. Lu H, Feng Y, Hu Y, et al. Spondin 2 promotes the proliferation, migration and invasion of gastric cancer cells. J Cell Mol Med. 2020;24(1):98–113. doi:10.1111/jcmm.14618

49. Liao CH, Yeh SC, Huang YH, et al. Positive regulation of spondin 2 by thyroid hormone is associated with cell migration and invasion. Endocr Relat Cancer. 2010;17(1):99–111. doi:10.1677/erc-09-0050

50. Ni H, Ni T, Feng J, Bian T, Liu Y, Zhang J. Spondin-2 is a novel diagnostic biomarker for laryngeal squamous cell carcinoma. Pathol Res Pract. 2019;215(2):286–291. doi:10.1016/J.PRP.2018.11.017

51. Su S, Chen J, Yao H, et al. CD10+GPR77+ Cancer-Associated Fibroblasts Promote Cancer Formation and Chemoresistance by Sustaining Cancer Stemness. Cell. 2018;172(4):841-856.e16. doi:10.1016/J.CELL.2018.01.009

52. LeBleu VS, Kalluri R. A peek into cancer-associated fibroblasts: origins, functions and translational impact. Dis Model Mech. 2018;11(4). doi:10.1242/DMM.029447

53. Sahai E, Astsaturov I, Cukierman E, et al. A framework for advancing our understanding of cancer-associated fibroblasts. Nat Rev Cancer 2020 203. 2020;20(3):174–186. doi:10.1038/s41568-019-0238-1

54. Li X, Balagam R, He TF, Lee PP, Igoshin OA, Levine H. On the mechanism of longrange orientational order of fibroblasts. Proc Natl Acad Sci U S A. 2017;114(34):8974–8979. doi:10.1073/PNAS.1707210114/SUPPL_FILE/PNAS.1707210114.SAPP.PDF

55. Jia W, Li H, He YW. The extracellular matrix protein mindin serves as an integrin ligand and is critical for inflammatory cell recruitment. Blood. 2005;106(12):3854–3859. doi:10.1182/blood-2005-04-1658

56. Bhattacharjee S, Hamberger F, Ravichandra A, et al. Tumor restriction by type I collagen opposes tumor-promoting effects of cancer-associated fibroblasts. J Clin Invest. 2021;131(11). doi:10.1172/JCI146987

57. Bhatt T, Dey R, Hegde AM, et al. The initiation of the wound healing program is regulated by the convergence of mechanical and epigenetic cues. bioRxiv. Published online October 22, 2021:2021.10.09.463764. doi:10.1101/2021.10.09.463764

58. Schindelin J, Arganda-Carreras I, Frise E, et al. Fiji: an open-source platform for biological-image analysis. Nat Methods 2012 97. 2012;9(7):676–682. doi:10.1038/nmeth.2019

59. Huang DW, Sherman BT, Lempicki RA. Systematic and integrative analysis of large gene lists using DAVID bioinformatics resources. Nat Protoc. 2009;4(1):44–57. doi:10.1038/NPROT.2008.211

60. Huang DW, Sherman BT, Tan Q, et al. The DAVID Gene Functional Classification Tool: a novel biological module-centric algorithm to functionally analyze large gene lists. Genome Biol. 2007;8(9):R183. doi:10.1186/GB-2007-8-9-R183

61. Huang DW, Sherman BT, Tan Q, et al. DAVID Bioinformatics Resources: expanded annotation database and novel algorithms to better extract biology from large gene lists. Nucleic Acids Res. 2007;35(suppl_2):W169–W175. doi:10.1093/NAR/GKM415

62. Carbon S, Douglass E, Good BM, et al. The Gene Ontology resource: enriching a GOld mine. Nucleic Acids Res. 2021;49(D1):D325. doi:10.1093/NAR/GKAA1113

63. Ashburner M, Ball CA, Blake JA, et al. Gene Ontology: tool for the unification of biology. Nat Genet. 2000;25(1):25. doi:10.1038/75556

64. Kanehisa M, Goto S. KEGG: Kyoto Encyclopedia of Genes and Genomes. Nucleic Acids Res. 2000;28(1):27. doi:10.1093/NAR/28.1.27

65. Piñero J, Ramírez-Anguita JM, Saüch-Pitarch J, et al. The DisGeNET knowledge platform for disease genomics: 2019 update. Nucleic Acids Res. 2020;48(D1):D845-D855. doi:10.1093/NAR/GKZ1021

