## Supplementary Files for "Mindin differentially regulates fibroblast subpopulations via distinct members of the Src family kinases during fibrogenesis"

**Supplementary Figure S1:**

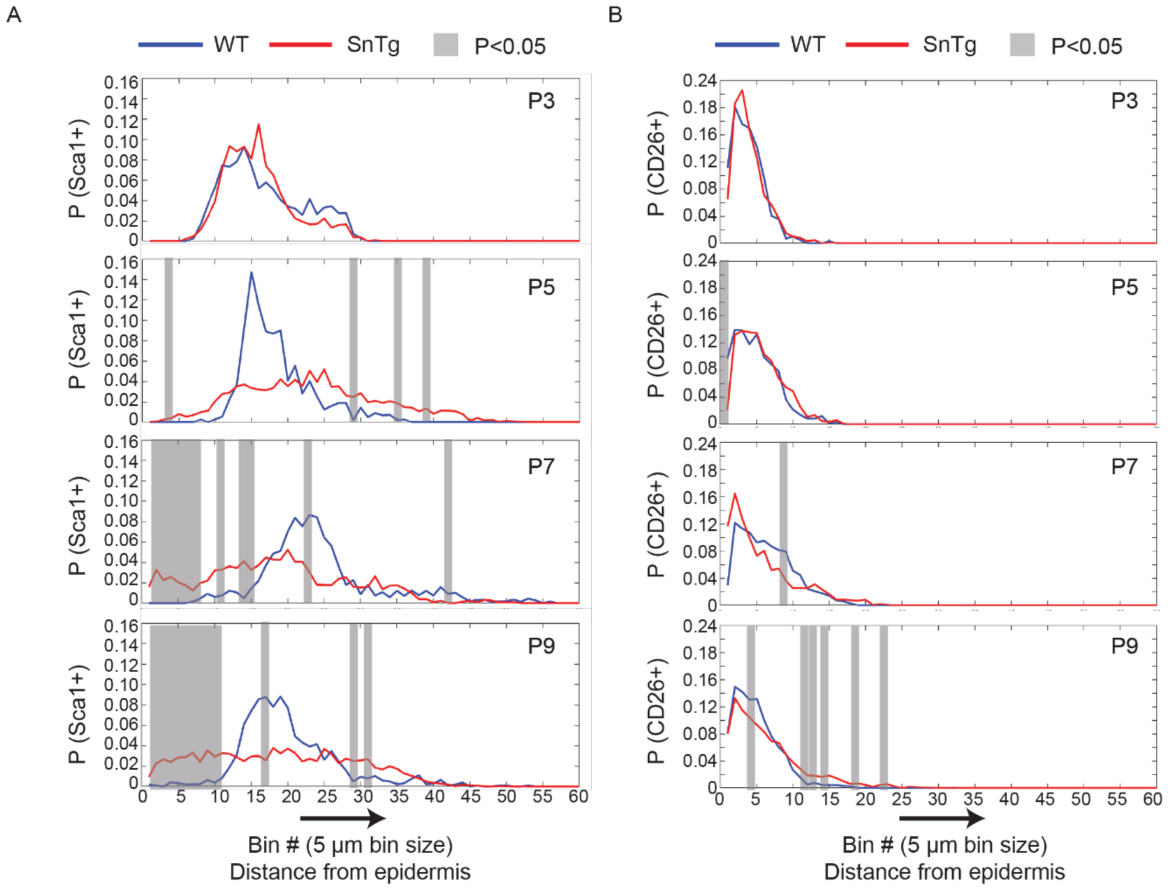


**Figure S1: Spatial distribution of Sca1+ dermal fibroblasts shift towards epidermis in SnTg skin.**
(A-B) Plots of the spatial probability distribution of (A) Sca1+ and (B) CD26+ cells in WT and SnTg skin taken from P3, P5, P7, and P9 pups. The x-axis corresponds to successive 5 µm thick bins starting from the epidermis towards the dermis. ($n \geq 3$, gray bars represent bins where $p<0.05$, using Welch's t-test).

**Supplementary Figure S2:**

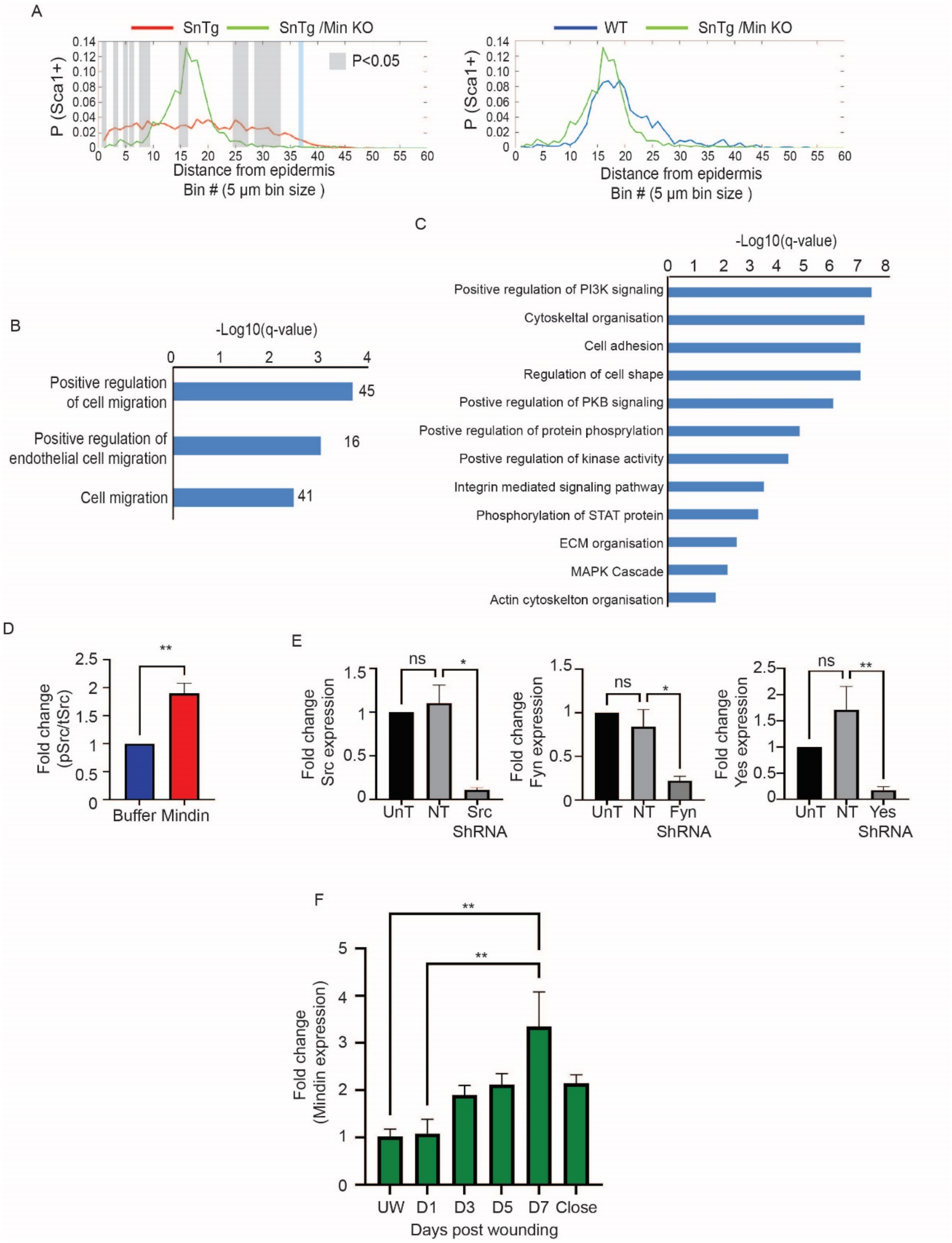


**Figure S2. Analysis of Mindin-induced migration in Sca1+ fibroblasts.** (A) Comparison of the spatial probability distribution of Sca1+ cells between SnTg and SnTg/Min KO (Top panel) and WT and SnTg/Min KO (bottom panel). Each successive bin# on the x-axis corresponds to successive 5 µm steps below the epidermis. ($n \geq 4$, area shaded in gray represent bins where $p<0.05$, calculated using Welch's t-test on corresponding bins). WT and SnTg are the same data as in Figure 1. (B) Gene set enrichment analysis (GSEA) showing biological processes associated with migration, using the differentially upregulated genes from Mindin treated fibroblasts. (C) GSEA of sub-list of Mindin upregulated genes associated with cell migration and positive regulation of cell migration. (D) Quantification of western blot for pSrc/tSrc w.r.t. to buffer control (n=4). (E) qPCR for comparison of RNA expression for Src, Fyn and Yes kinases in the cells transduced with Non-targeting (NT) shRNA and Src, Fyn or Yes shRNA respectively (n≥3). (F) qPCR for Mindin expression at 3-, 5-, 7- and 10-days post wounding (dpw) relative to unwounded skin (n≥3 mice for each group). Wounds closed on 10dpw. Data represents the mean ± SEM. p-values were calculated by Ratio paired t-test (E), 1-way ANOVA followed by Tukey's post hoc analysis (F and G) (*P < 0.05, **P < 0.01, ***P < 0.001, ****P < 0.0001 and ns (P>0.05) is non-significant).

**Supplementary Figure S3:**

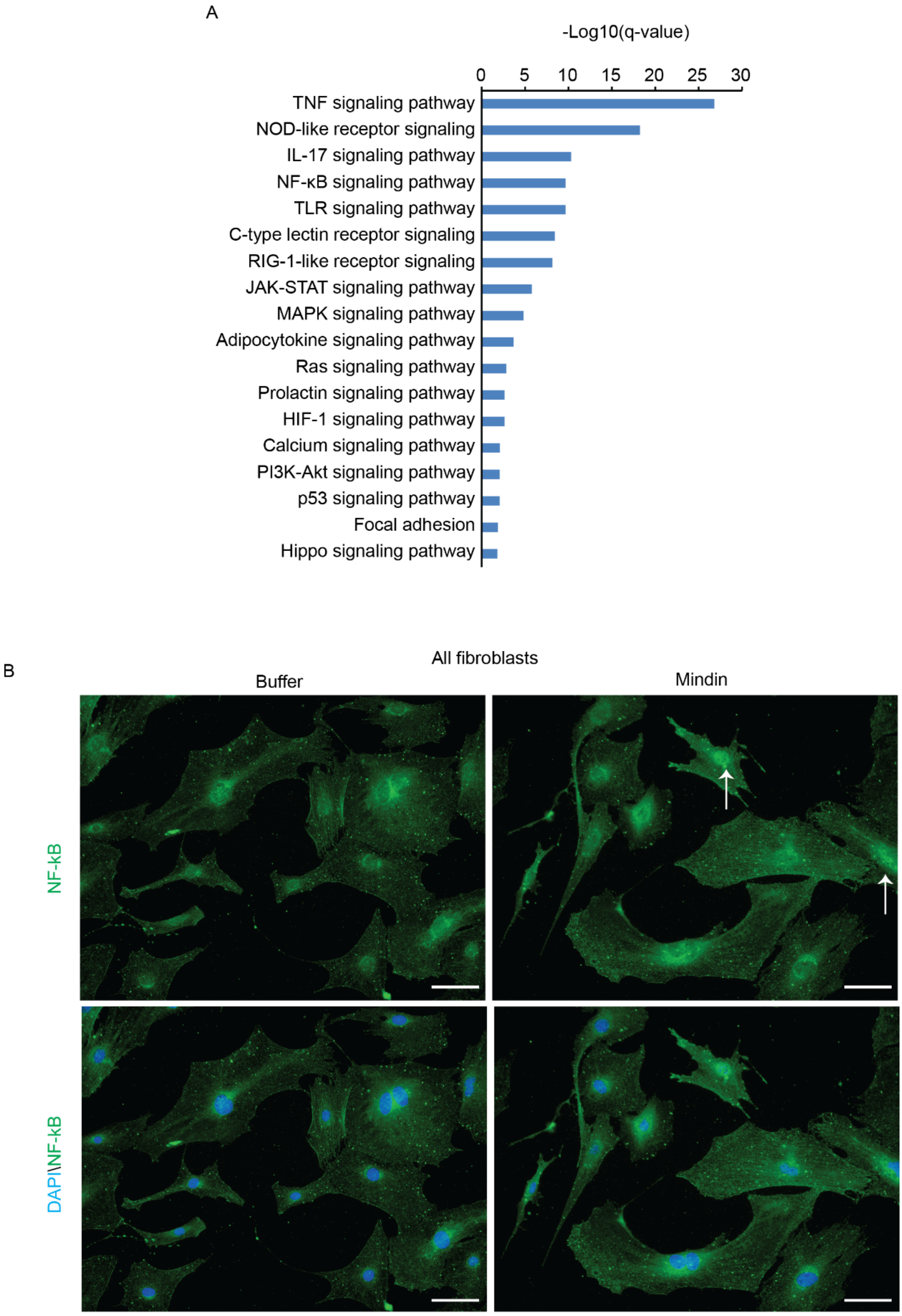


**Figure S3: Mindin activates inflammatory pathways in fibroblasts** (A) Enriched KEGG pathways terms based on GSEA of genes upregulated (1715 genes, FC>1.5, q<0.05) upon treatment of fibroblasts with Mindin. (B) IF staining for NF-κB (green) and DAPI (blue) in unsorted mixed fibroblasts treated with either buffer or Mindin. White arrows point to +ively stained nucleus (magnification 20x, scale bar = 50 μm).

**Supplementary Figure S4:**

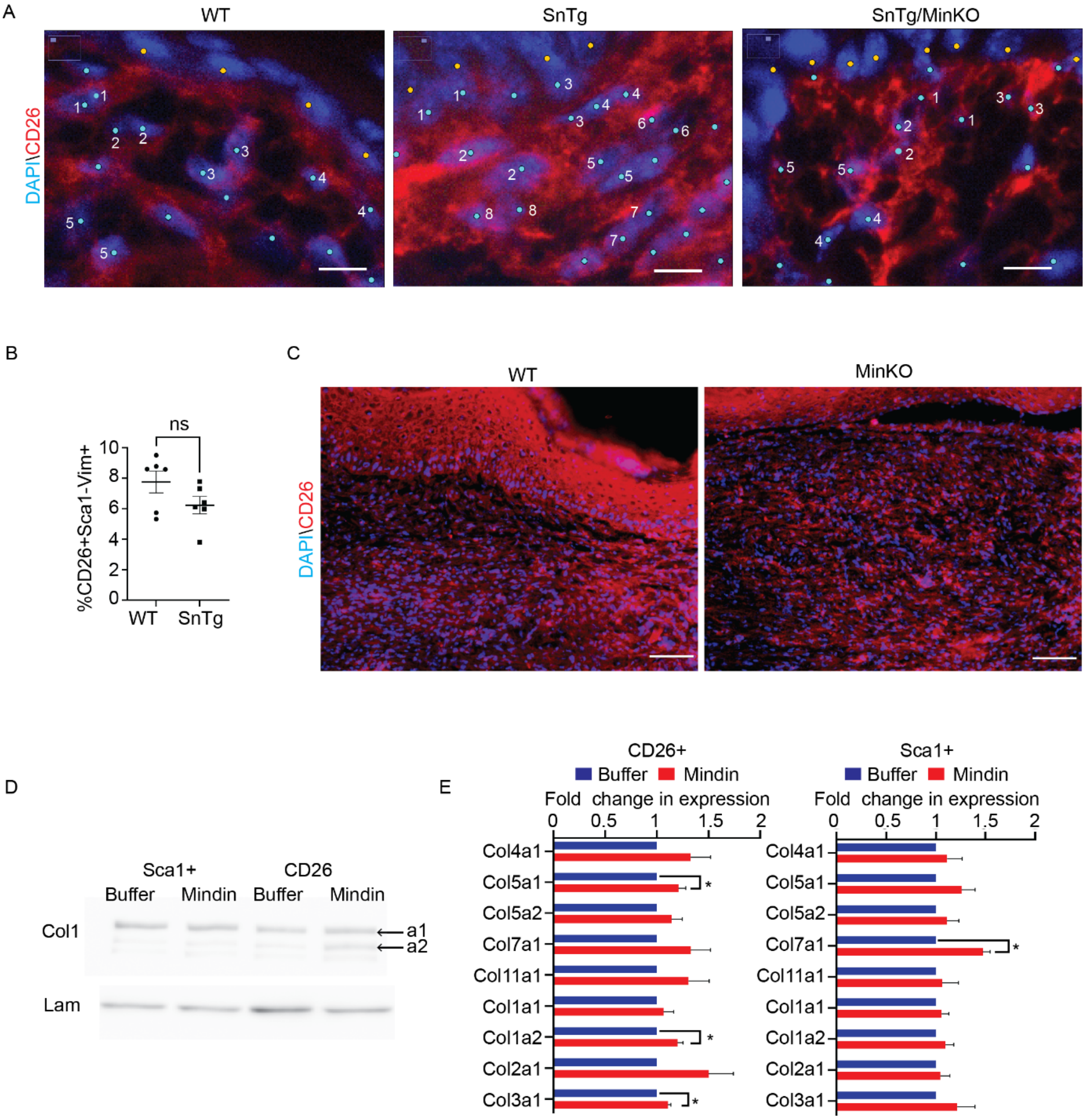


**Figure S4. CD26+ fibroblasts contract and produce collagen in response to Mindin.** (A) IF of CD26 (Red) and DAPI (blue) in WT, SnTg, and SnTg Min KO skin sections (scale bar = 5 µm). Numbers refer to representative neighbouring cell pairs that are used to calculate intercellular distance between nearest neighbours. (B) Percentage of CD26+Sca1-Vim+ cells in WT and SnTg mice, measured via FACS. (each dot represents an individual animal) (C) IF of CD26 (Red) and DAPI (blue) in WT and Min KO in wounded skin sections 7 days post-wounding (scale bar = 50 µm). (D) Western blot showing bands for Collagen 1 (Col1) and Lamin B1 in Sca1+ and CD26+ fibroblasts treated with either buffer control or Mindin. (E) RNA expression of different collagen subtypes quantified by RT-PCR in Mindin or buffer treated CD26+ and Sca1+ fibroblasts (n = 4). Data represents the mean ± SEM. p-values were calculated by Welch's t-test (B), ratio-paired t-test (E) (*P < 0.05 and ns (P>0.05) is non-significant).

**Supplementary Figure S5:**

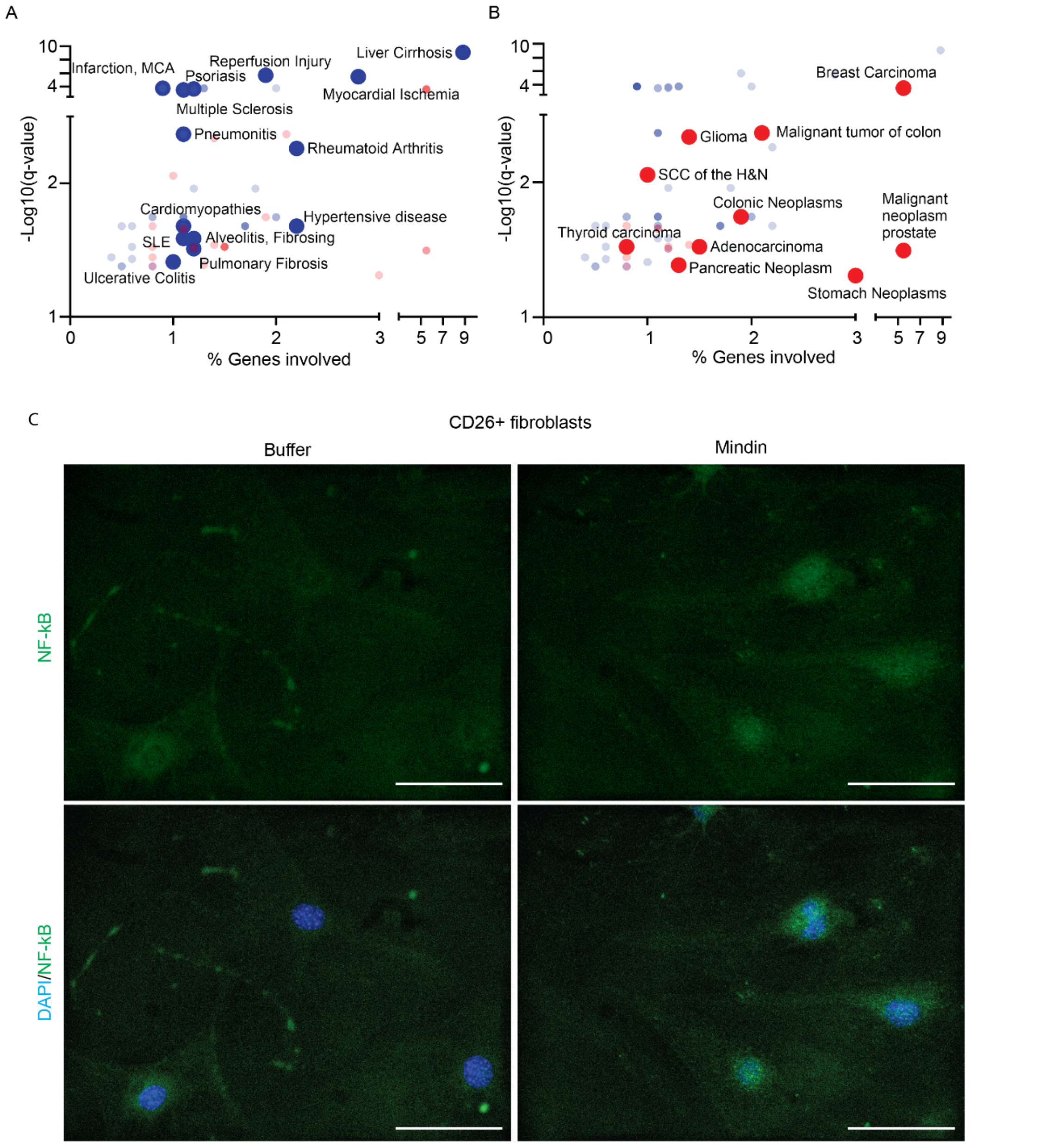


**Figure S5:** (A-B) GSEA of genes upregulated in Mindin treated fibroblasts, revealed terms associated with (A) fibrotic and inflammatory diseases and (B) cancers (using the DisGeNET database). (Plotted as % genes of total upregulated genes (1715) in the list (x-axis) and -Log10(q-value) (y-axis). (C) IF staining for NF-kB (green) and DAPI (blue) in CD26+ fibroblasts treated with either buffer (left) or Mindin (right) for 24-hours.

**Supplementary Figure S6:**

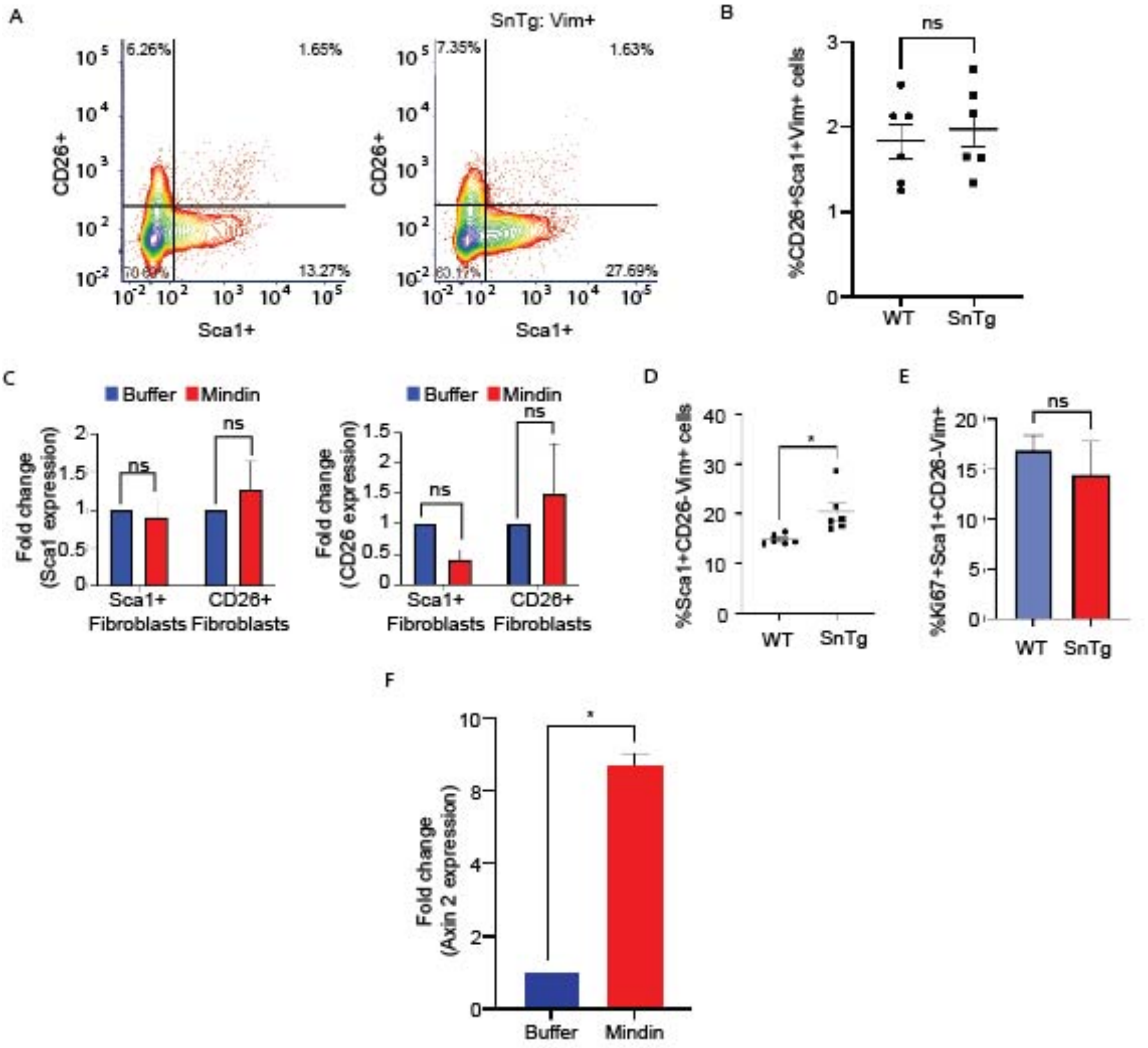


**Figure S6: Sca1+ fibroblasts are increased in SnTg skin.** (A) Representative contour plots showing Vim+ cells gated for CD26 and Sca1. (B) %CD26+Sca1+Vim+ cells in WT and SnTg skin (each dot represents individual mice; n=6). Fold change in RNA expression of Sca1 (left panel) and CD26 (right panel) in CD26+ and Sca1+ fibroblasts treated with either buffer of Mindin. (D) %Sca1+CD26-Vim+ cells in WT and SnTg skin (each dot represents individual mice; n=6). (E) %Ki67+ of Sca1+CD26-Vim+ cells in WT and SnTg skin (n=3). (F) Fold change in RNA expression level of Axin2 in Sca1+ fibroblasts treated with either buffer of Mindin (n=2). Data represents the mean ± SEM. p-values were calculated by Welch's t-test (B,D,E), or ratio paired t-test (C,F), (*P < 0.05, ns (P>0.05) is non-significant).

| Supplementary Table 1 | | |
|---|---|---|
| Gene | Forward primer | Reverse Primer |
| GAPDH | AGGTCGGTGTGAACGGATTTG | TGTAGACCATGTAGTTGAGGTCA |
| β-Actin | GGGCTATGCTCTCCCTCAC | GATGTCACGCACGATTTCC |
| Col1a1 | GCCAAGAAGACATCCCTGAAG | TCATTGCATTGCACGTCATC |
| Col1a2 | TGCTGCTTGCAGTAACGTCG | TCAACACCATCTCTGCCTCG |
| Col2a1 | GGGAATGTCCTCTGCGATGAC | GAAGGGGATCTCGGGGTTG |
| Col3a1 | CTGTAACATGGAAACTGGGGAAA | CCATAGCTGAACTGAAAACCACC |
| Col4a1 | CCTGGCACAAAAGGGACGA | ACGTGGCCGAGAATTTCACC |
| Col5a1 | GCCCTCAGGGGTAACGAAAAC | GACTCGGTAGGCAACATCCG |
| Col5a2 | TTGGAAACCTTCTCCATGTCAGA | TCCCCAGTGGGTGTTATAGGA |
| Col7a1 | ACCACGTTTCTGACCGTGTC | AGCTGTGTCCACTAAATCTTGG |
| COL11a1 | CCAGCGGGTCTTATGGGTC | TGGTAACATCAGCATGGTTCC |
| Rantes | CCTCACCATCATCCTCACTGCA | TCTTCTCTGGGTTGGCACACAC |
| IL6 | CGTGGAAATGAGAAAAGAGTTGTG | CCAGTTTGGTAGCATCCATCATTTCT |
| CXCL10 | CCAAGTGCTGCCGTCATTTTC | GGCTCGCAGGGATGATTTCAA |
| CXCL5 | TGCGTTGTGTTTGCTTAACCG | CTTCCACCGTAGGGCACTG |
| CXCL3 | CTGCACCCAGACAGAAGTCAT | CCGTTGGGATGGATCGCTTT |
| IL17 | GGAGAGCTTCATCTGTGTCTCTG | TTGGCCTCAGTGTTTGGACA |
| Sca1 | AGGAGGCAGCAGTTATTGTGG | CGTTGACCTTAGTACCCAGGA |
| CD26 | ACCGTGGAAGGTTCTTCTGG | CACAAAGAGTAGGACTTGACCC |
| Axin2 | TGACTCTCCTTCCAGATCCCA | TGCCCACACTAGGCTGACA |
